# Transcriptome profiling of *Candidatus* Liberibacter asiaticus in citrus and psyllids

**DOI:** 10.1101/2021.08.09.455679

**Authors:** Agustina De Francesco, Amelia H. Lovelace, Dipan Shaw, Min Qiu, Yuanchao Wang, Fatta Gurung, Veronica Ancona, Chunxia Wang, Amit Levy, Tao Jiang, Wenbo Ma

## Abstract

*Candidatus* Liberibacter asiaticus (Las) is an emergent bacterial pathogen that is associated with the devastating citrus Huanglongbing (HLB). Vectored by the Asian citrus psyllid, Las colonizes the phloem tissue of citrus, causing severe damage to infected trees. So far, cultivating pure Las culture in axenic media has not been successful and dual-transcriptome analyses aiming to profile gene expression in both Las and its host(s) have a low coverage of the Las genome due to the low abundance of bacterial RNA in total RNA extracts from infected tissues. Therefore, the lack of a Las transcriptome remains as a significant knowledge gap. Here, we used a bacterial cell enrichment procedure and confidently determined the expression profiles of approximately 84% of the Las genes. Genes that exhibited the highest expression levels in citrus include ion transporters, ferritin, outer membrane porins, and genes involved in phage-related functions, pilus formation, cell wall modification, and stress responses. One hundred and six genes were found to be differentially expressed in citrus vs psyllids. Genes related to transcription/translation and resilience to host defense response were upregulated in citrus; whereas genes involved in energy generation and the flagella system were expressed to higher levels in psyllids. We also determined the relative expression levels of potential Sec-dependent effectors, which are considered as key virulence factors of Las. This work advances our understanding of HLB biology and offers novel insight into the interactions of Las with its plant host and insect vector.

## Introduction

Citrus Huanglongbing (HLB) was first described in China in 1919. Over the years, HLB has been reported from all citrus-growing areas worldwide and is now considered to be the most devastating disease of citrus (Bové 2006; Wang and Trivedi 2013). After the introduction of HLB in Florida, in 2005 (Gottwald 2010), citrus production has decreased by 70%, causing billions of dollars of economic losses (Hodges and Spreen 2012; Singerman and Rogers 2020). Since then, HLB has spread to other citrus-producing states in the US, especially Texas and California, posing an unprecedented threat to the industry (da Graça et al. 2016; Graham et al. 2020).

HLB damages citrus production by decreasing the lifespan of the infected trees, reducing fruit yield, and affecting the fruit quality (Bové 2006; Graham et al. 2013). Typical disease symptoms include blotchy mottling and yellowing of leaves; canopy defoliation and dieback; small, deformed, unevenly-colored, and off-tasting fruits; premature fruit drop; and loss of fibrous roots and overall root decline, ultimately leading to tree death (Bové 2006; Gabriel et al. 2020). Pathogens associated with HLB are species of the Gram-negative bacteria *Candidatus* Liberibacter. Transmitted by insect vectors, Liberibacter colonizes the sieve elements of the phloem tissue in citrus trees. Despite the devastating impact of HLB on citrus industry, no curative treatment or el□ective management methods are currently available to eliminate or mitigate Liberibacter in infected trees (Munir et al., 2018) and management of HLB has focused on preventing the spread of the disease by removing infected trees and controlling insect vectors (Grafton-Cardwell et al. 2013). Thus, there is an urgent need to explore new control strategies, which requires a better understanding of the basic biology and virulence mechanisms of Liberibacter (Wang et al. 2017; Wang 2019). Unfortunately, this research has been hindered by the inability to grow an axenic culture of pathogenic Liberibacter despite enormous efforts over the past years. As a result, biological functions and pathways that are important for its colonization in citrus are poorly understood.

Three Liberibacter species have been found to associate with HLB: Liberibacter asiaticus (Las), Liberibacter africanus (Laf), and Liberibacter americanus (Lam) (Bové 2006). Las and Lam are transmitted by the Asian citrus psyllid (ACP, *Diaphorina citri*), while Laf is transmitted by the African citrus psyllid (*Trioza erytreae*) (Wang et al. 2017). Las is the most widely-distributed species and is predominantly associated with HLB in Asia, Africa, the Middle East, and the Americas (Gottwald 2010; Wang 2019). Due to its low and uneven titer in infected citrus, the first genome of Las was obtained using infected insects (Duan et al. 2009). Recently, genome sequences of several isolates from different geographic areas were determined, providing insights into the phylogenetic relationship and genomic variation of Las (Thapa et al. 2020).

Several Las genes have been studied for their potential roles in host interaction and disease development. Flagellin is a well-studied bacterial feature that is often recognized by the plant innate immune system to activate defense response. Although Las harbors genes encoding the flagella system, flagella are only formed in ACP but not in citrus (Andrade et al. 2020). Another gene that may manipulate plant immunity encodes a salicylic acid (SA) hydroxylase, which hydrolyzes the plant defense hormone SA and thus inhibits defense response (Li et al. 2017). Furthermore, a peroxidase encoded by the prophage SC2 may function as a secreted effector and help detoxify reactive oxygen species (ROS) produced by the host as a defense mechanism (Jain et al. 2015). Finally, proteins secreted through the Sec secretion system have been investigated for their role in facilitating Las colonization in citrus. A core set of Sec-dependent effectors (SDEs) have been identified from Las isolates (Pagliaccia et al. 2017; Thapa et al. 2020) and the virulence activity of some of these SDEs has been characterized (Clark et al. 2018; Liu et al. 2019; Clark et al. 2020; Pang et al. 2020).

Although transcriptome analysis is widely, and routinely, used to understand biological processes, assessment of the Las transcriptome has been challenging due to the extremely small proportion of bacterial transcripts in total RNA extracted from infected tissues. In fact, thorough analysis of bacterial transcriptome *in planta* has only recently been accomplished for the model pathogen *Pseudomonas syringae* using protocols that either enrich bacterial cells before RNA extraction (Nobori et al. 2018) or enrich bacterial RNAs in total RNA extract (Lovelace et al. 2018). To date, transcriptome studies of HLB have focused on host responses by comparing gene expression changes in healthy vs infected citrus or ACP (e.g Kruse et al. 2017; Liu et al. 2020; Wei et al. 2021) and in different citrus varieties with and without infection (e.g. Wang et al. 2016; Rawat et al. 2017; Hu et al. 2017; Fang et al. 2021; Chin et al. 2021). However, the Las transcriptomic landscape has not been determined, representing a significant knowledge gap. Here, we evaluated various workflows to optimize RNA-seq outputs for Las using infected citrus and ACP tissues. Our best performing protocol allowed the determination of expression levels for 84% Las genes in citrus and 100% in psyllids and facilitated the subsequent analysis of differentially expressed genes in citrus vs psyllids. These analyses provide new information on the cellular processes in Las that might be important for their interactions with the plant host and/or insect vector.

## Materials and Methods

### Plant and insect materials

Citrus and ACP samples were obtained from fields in Florida and Texas that have been heavily exposed to and affected by HLB. Healthy samples of both hosts were taken as controls. Midrib tissues were collected from two citrus varieties: Rio Red grapefruit (*Citrus paradisi* Macfadyenand) and *Citrus macrophylla*. Four biological replicates of HLB-infected Rio Red grapefruit were taken from symptomatic, HLB-infected trees that were at least seven years-old in a field in Texas. Las-positive *C. macrophylla* and insects (5 biological replicates from different colonies) were collected in Florida. Las infection was confirmed and their titer was determined by reverse transcription-quantitative polymerase chain reaction (RT-qPCR) using Las 16S primers (Yan et al. 2013).

### Bacterial isolation from infected citrus tissue

This pre-enrichment procedure was applied to citrus samples using a protocol modified from Nobori et al. (2018). Freeze-dried midribs (from 15–25 leaves) were ground to a fine powder in liquid nitrogen. 30 mL of fresh bacterial isolation buffer (9.5% ethanol 0.5% phenol 2% β-mercapto-ethanol pH 4.5) was added and mixed thoroughly by vortex. All subsequent processes were done on ice or in a cold room (4°C). The samples were incubated for 20 h with slow shaking, and then passed through a Miracloth® (EMD Millipore) filter to remove large plant debris. The flow through was centrifuged at 3,200 × *g* for 30 min to pellet plant and bacterial cells. The supernatant was carefully removed, and the pellet was resuspended in 900 μL of a buffer containing 9.5% EtOH and 0.5% phenol. The suspension was centrifuged at 2,300 × *g* for 20 min to obtain a two-layered pellet (a white layer on the top and a green layer on the bottom). The top-white layer (bacterial cells) was resuspended by pipetting while keeping the bottom layer (plant cells) undisturbed. This step was repeated 2-3 times until obtaining a pure white layer. Bacterial cells were collected by centrifuging at 10,000 × *g* for 2 min and resuspended in 1 mL of TLE buffer (200 mM Tris-HCl (pH 8.2), 100 mM LiCl, 50 mM EDTA, 10% SDS, 2% β-mercaptoethanol) for RNA extraction.

### RNA extraction

For RNA extraction from the citrus midrib, 1 mL of TLE buffer was added to 100 mg citrus powder. 60 μL of 500 mM CaCl_2_ was added to the suspension or the bacterial cells isolated from the citrus tissue as described above, mixed thoroughly, and placed on ice for 10 min. The solution was centrifuged at 12,000 × *g* for 10 min at 4°C, followed by two rounds of extraction using 500 μL of phenol:chloroform:isoamyl alcohol (25:25:1) and centrifugation at 15,000 × *g* for 10 min at 4°C. The supernatant was mixed with 2 M LiCl and incubated overnight at −20°C. RNA was precipitated by centrifugation at 15,000 × *g* for 30 min, washed twice with ice-cold 75% ethanol, and re-suspended in 20 μL of RNAse-free water.

For RNA extraction from ACP, 10-15 adult psyllids were ground in liquid nitrogen, immediately suspended in 50 μL TRIzol® (Invitrogen) and ground until complete homogenization. An additional 950 μL of TRIzol® was added to the homogenate, followed by extraction using 200 μL of chloroform and final precipitation using 500 μL of isopropanol. The RNA pellet was washed twice using 75% ethanol and resuspended in 20-30 μL of RNAse-free water.

RNA extracts were treated with DNase I (Thermo-Fisher Scientific). The RNA concentration and quality of the extracts were determined by nanodrop and an Agilent 2100 Bioanalyzer using the eukaryotic analysis software.

### Enrichment of Las RNA

Eukaryotic RNA (18S and 28S rRNAs and polyadenylated RNAs) was removed from total RNA (10-25 μg) extracted from infected tissues by MICROBEnrich™ kit (Invitrogen) following manufacturer’s instructions. Some of the RNA samples were further treated by using the RiboMinus™ Eukaryote Kit (Invitrogen) in order to completely deplete 5S and 5.8S rRNA as well as residual 18S and 28S rRNA. Some RNA samples, following the MICROBEnrich + RiboMinus™ Eukaryote Kit treatments were further treated with Poly(A) Polymerase (NEB), which adds a Poly(A) tail to bacterial mRNA. The resulting RNA samples were then proceeded for library construction and RNA-seq.

### Quantitative reverse transcription PCR (RT-qPCR)

We examined the effectiveness of the bacterial RNA enrichment following various treatments by comparing the expression levels of Las 16S rRNA gene (Yan et al. 2013) and F-Box gene (Citrus Unigene ID CAS-PT-306416) from *Poncirus trifoliata* (Mafra et al. 2012) using RT-qPCR. Primer sequences are listed in Supplemental Table 1. cDNA was synthesized from 1 μg of total DNase-treated RNA using 300 μM of random primers (Invitrogen) and the RevertAid First Strand cDNA Synthesis Kit (Thermo Scientific) in 20 μL final volume. qPCR reactions were performed in a CFX96 real-time PCR detection system (Bio-Rad) containing 100 ng of cDNA template (2 μL), 2X Quantitect SYBR Green (BioRad), RNase-free water and 10 μM of gene-specific primers in a final volume of 10 μL. Cycling conditions were: 95°C for 2 min, followed by amplification with 43 cycles of 95°C for 15 sec, 54°C for 30 sec, and 72°C for 30 sec. Relative abundance of the Las RNA was calculated using the equation ΔCt = Ct _Las 16s_ − Ct _citrus Fbox_ where the Ct values were the average of two technical replicates.

RT-qPCR for Sec-dependent effector (SDE) candidates in citrus was performed using the same procedure described above and primers specified in Supplemental Table 1. The one-step QuantiTect SYBR Green RT-PCR kit (Qiagen) was used to synthesize the cDNA and perform qPCR under the following conditions: 50°C for 30 min, 95°C for 15 min, and then 40 cycles of 94°C for 10 sec, 58°C for 30 sec, and 72°C for 30 sec. SDE expression was confirmed when Las-positive samples gave a positive Ct value and healthy controls did not allow amplification. For some genes, the primers allowed a minimal amplification using RNA extracted from healthy citrus tissues (confirmed by qPCR using DNA extract and 16S rDNA primers) and gave a high C value. In these cases, ΔCt was calculated as Ct _Las gene in healthy citrus_ − Ct _Las gene in diseased citrus_ using the average values of technical duplicates and at least two biological replicates. A t-test for pairwise comparison (95% confidence) was used to determine whether expression of the SDE was significant.

**Table 1.** Comparison of various bacterial cell or RNA enrichment protocols using Las-infected citrus midrib tissues.

| Sample<sup>a</sup> | Las titer<sup>b</sup> | Treatment<sup>c</sup> | Relative abundance of Las 16S rRNA<sup>d</sup> |
| --- | --- | --- | --- |
| Macrophylla | 22.82 | none | -1.70 |
|  |  | 1-step | 0.01 |
|  |  | 2-step | 1.78 |
| Grapefruit 1 | 23.23 | none | -0.94 |
|  |  | 1-step | 0.16 |
|  |  | 2-step | 1.87 |
| Grapefruit 2 | 20.67 | none | -5.69 |
|  |  | bacterial isolation | -6.41 |
| Grapefruit 3 | 22.44 | none | -1.93 |
|  |  | bacterial isolation | -3.24 |
| Grapefruit 4 | 22.67 | none | -1.49 |
|  |  | bacterial isolation | -2.28 |

<sup>a</sup> Citrus host and biological replicate for HLB-infected sample.
<sup>b</sup> Ct value of Las 16S rRNA determined by qRT-PCR.
<sup>c</sup> no enrichment (none), 1- or 2-step treatments were done using total RNA extract, and bacterial isolation was a treatment before RNA extraction.
<sup>d</sup> Relative abundance of Las to citrus genes was calculated as $\Delta Ct = (Ct_{\text{Las 16S}} - Ct_{\text{citrus Fbox}})$ . Negative values indicate higher relative abundance of Las.

### RNA-seq

Most of the sequencing-ready libraries were constructed using TruSeq Stranded Total RNA Library Prep Kit (Illumina). The RNA samples were treated using the Illumina Ribo-Zero Plus rRNA depletion kit as part of the standard protocol. For the RNA samples treated with Poly(A) Polymerase, Poly(A)-enriched library construction was utilized. cDNA libraries were then subjected to RNA-seq using an Illumina HiSeq-4000 system with 150-bp strand-specific single-end reads and technical duplicates. Library preparation and sequencing were performed in the DNA Technologies Core Genome Center at University of California, Davis. The total reads per library yielded in these experiments ranged from 83 to 848 million reads.

### RNA-seq analysis

The quality of the raw sequencing data was checked with FastQC v0.11.9 (Andrews 2010). Subsequent statistical analyses were performed using R and Python. To ensure high-quality sequences for mapping and downstream analyses, the reads were trimmed using Trimmomatic v 0.36. The Trimmomatic program comes with standard adapter files (Bolger et al. 2014). RNA-seq reads were aligned with the indexed Las-Psy62 genome assembly (NC_012985.3) and parsed to it using HISAT2 v2.1.0. The number of reads mapped to each gene was counted using the feature counts function of the Rsubread package (Liao et al. 2019) and listed in Supplemental Data Set 1. All data used for analysis have been deposited in the GEO database (accession No. 180624).

### Differential Gene Expression analysis

Gene expression values were normalized to counts per million (CPM). Low-expression genes were filtered using the row sum function in R with cpm > 0.5 and rowSum >= 5 (Robinson et al. 2010). The retained genes included for differential expression had over 0.5 CPM in at least four samples (Robinson et al. 2010). Differentially expressed genes (DEGs) were identified from gene counts for each sample using the DESeq2 package v1.30.1 from Bioconductor (Love et al. 2014). DEGs were selected based on log_2_-transformed and normalized counts that have an adjusted *p* value (padj) below a false discovery rate (FDR) cutoff of 0.05. Principal components analysis (PCA) of rlog transformed read counts was performed for all treatments and replicates using the plotPCA function in DESeq2. The rlog function transforms the count data to the log_2_ scale in a way that minimizes differences between samples with low counts. Hierarchical clustering and Euclidean distance clustering were performed on normalized read counts between treatments and replicates using the dist function and pheatmap function of the pheatmap library v1.0.12.

### Pathway analysis

Gene set (or pathway) analysis was conducted using log_2_ fold change (log_2_FC) values of normalized mean counts for annotated genes using the Bioconductor package, gauge version 2.24.0. Gene sets of metabolic pathways were obtained from the KEGG pathway database (Kanehisa et al. 2008), using the organismal code “las” for Las psy62. Significant gene sets were identified from log_2_ fold changes between treatments and were selected based on an FDR q value cut-off of 0.05.

Relevant gene ontology (GO) terms were recruited manually from the KEGG gene database, where we determined the IDs of the associated proteins in Uniprot DataBase and extracted their complete GO annotations from QuickGO version 2021-02-08. The GO-term analysis was conducted using the biocManager package, ViSEAGO v.1.4.0. The custom GO-terms for Las were annotated using the Custom2GO and its annotation function. Enriched GO-terms were identified for each cluster and DEG set using the entire genome as the background gene set using the “create_topGO data” function. To test for significant enrichment, the classic algorithm and fisher exact test were used with a cutoff *p* value of 0.05. Redundant enriched GO terms were identified using the REVIGO web server (Supek et al. 2011).

### Correlation analysis of the RNA-seq data with published RT-qPCR data

Correlation analysis was conducted on log_2_FC values of Las genes found in citrus midribs relative to those found in ACP samples using the cor.test function in the R stats package v. 4.0.4. A total of 225 Las genes with differential relative expression in citrus midribs and ACP were previously evaluated by RT-qPCR by Yan et al. (2013) and Thapa et al. (2020) and were used to compare to our RNA-seq data set. The entire gene set from each published data set as well as gene sets from representative pathways defined by Yan et al. (2013) were evaluated for normal distribution using the shapiro.test function. Correlation was evaluated using Pearson’s correlation test on the normally distributed data from Thapa et al. (2020), whereas the Spearman’s test was used on the non-normally distributed data from Yan et al. (2013). For both tests, the significance was evaluated using a *p* value cutoff of 0.05.

## Results

### Enrichment of Las RNA from infected citrus and ACP tissues

Our understanding of Las biology is hindered by the lack of knowledge of its transcriptome landscape. To overcome this limitation, we first aimed to establish a protocol that enables improved coverage of the Las genome in RNA-seq experiments. Because Las RNA is vastly underrepresented in total RNA extracted from infected host tissues, especially in citrus, several bacterial RNA enrichment methods were explored (Fig. 1).

**Fig 1.**
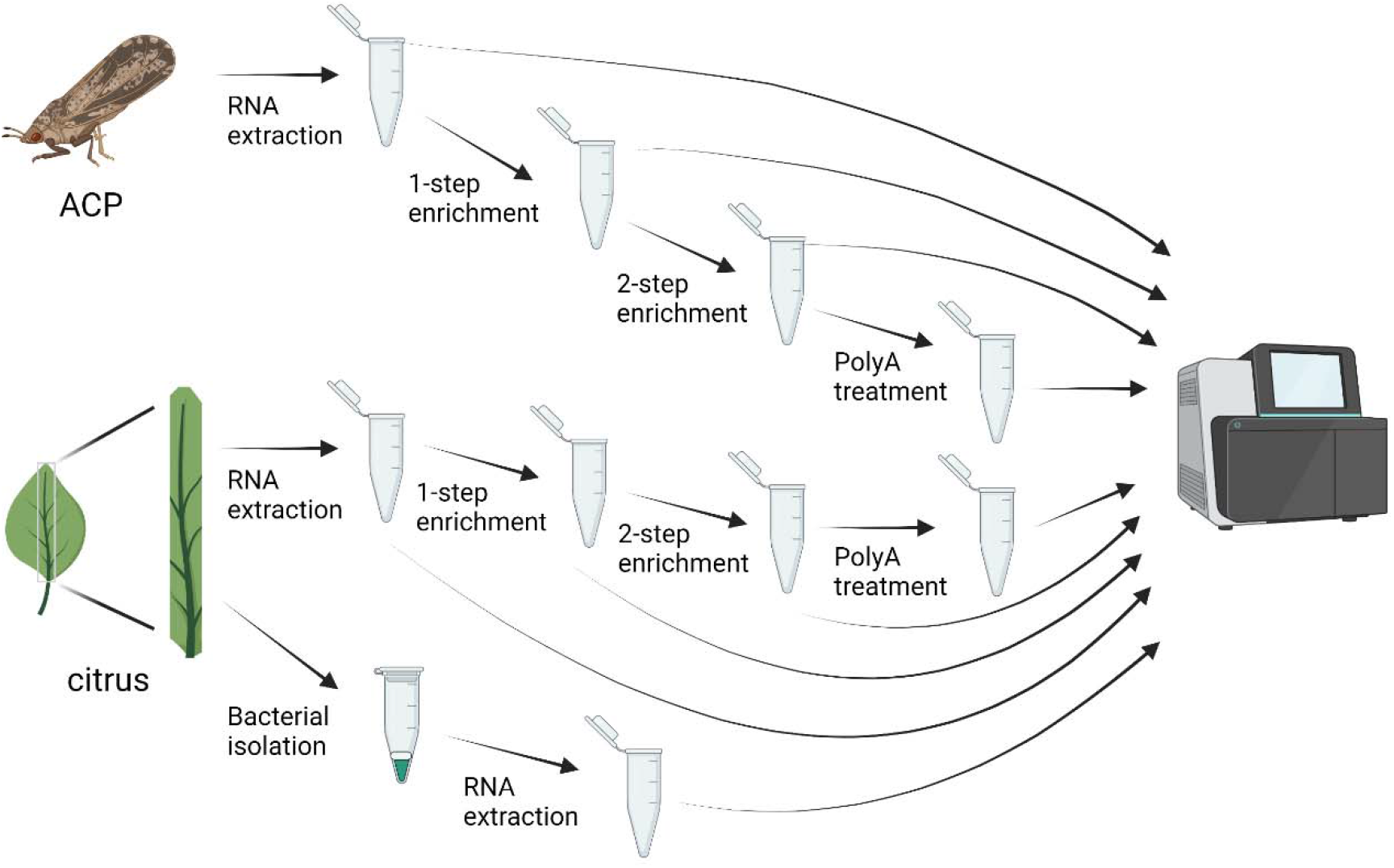
RNA sample preparation workflow. Scheme of the experimental procedures examined to increase the proportion of Las RNA in samples prepared from HLB-infected citrus and Asian citrus psyllids (ACP) tissues for RNA-seq. For RNA enrichment, total RNA extracts were subjected to treatment using MICROBEnrich™ (1-step) followed by an additional step using RiboMinus™ Eukaryote Kit (2-step). Some of the 2-step enriched RNA samples were further treated with Poly(A) Polymerase (PolyA) from which bacterial RNAs would be ligated to a poly(A) tail to facilitate Poly(A)-enriched RNA-seq library construction. The midrib tissues were also subjected to a bacterial isolation procedure to enrich Las cells before RNA extraction. All RNA samples were subjected to RNA-seq analyses with non-treated samples as the controls. Except for the PolyA treated samples, the RNA samples were further treated using Ribo-Zero Plus rRNA depletion kit as part of the standard RNA-seq library construction protocol. Illustration was made by Biorender.com.

For citrus tissues, midribs were separated from leaves of grapefruit (*Citrus paradisi* Macfadyenand) and macrophylla (*Citrus macrophylla*) trees with HLB symptoms. Las titer in each sample was determined by RT-qPCR using 16S primers (Table 1, Supplemental Table 1). Total RNA was extracted and subsequently subjected to the MICROBEnrich™ kit (Invitrogen) (1-step treatment) or a combination of MICROBEnrich™ and the RiboMinus™ Eukaryote Kit (Invitrogen) (2-step treatment) to deplete plant RNAs, which constitute the vast majority of the RNA. The effectiveness of Las RNA enrichment in the resulting RNA samples was evaluated using bioanalyzer and RT-qPCR in comparison to non-treated samples.

Electropherogram from the bioanalyzer shows a drastic decrease of 28S and 18S rRNA after the 1-step treatment (Supplemental Fig. 1). However, the amount of 16S rRNA, represented by absolute intensity (FU), also decreased after the 1-step treatment, representing a general loss of RNA or loss of 16S rRNA in addition to eukaryotic rRNA and mRNA. Additional depletion of both prokaryotic and eukaryotic rRNAs was observed after 2-step treatment as the rRNA peaks were no longer distinguishable (Supplemental Fig. 1). RT-qPCR results showed a consistent decrease in the relative abundance of Las 16S rRNA over a housekeeping gene of citrus (Table 1) after the successive treatments. This indicates that either these treatments were ineffective in enriching Las RNA, or 16S rRNA was not informative to indicate the relative abundance of bacterial RNA after the treatments.

In addition to enriching bacterial RNA from total RNA extract, we also enriched bacterial cells from homogenized midrib tissues using a centrifugation-based procedure (called bacterial isolation hereinafter). RT-qPCR results showed that the relative transcript abundances of Las 16S rRNA had consistent increases compared to the non-treated controls (Table 1), indicating successful enrichment of Las cells and a relatively higher proportion of Las RNA in the RNA extract.

### RNA-seq analysis using enriched Las RNA samples

All RNA samples described above were subjected to RNA-seq. In total, we performed RNA-seq using RNAs from one macrophylla sample, one grapefruit sample, three bacterial cell isolation samples (from grapefruits), and five ACP samples, with technical replicates. Among them, multiple libraries were sequenced for the macrophylla, grapefruit and two of the ACP samples using RNA that went through various enrichment treatments (Fig. 1).

Total RNA-seq reads were parsed to the genome assembly available for the psy62 strain of Las. Based on the alignment, the raw read counts corresponding to Las were calculated for each gene and the read depth represents the total number of reads aligned to the Las genome. A wide range of read depth values and detected Las genes was found among samples and treatments (Table 2, Fig. 2). In general, 1-step and 2-step enrichment marginally increased the number of reads mapped to Las genes and the coverage of the Las genome; 2-step (PolyA) treatment actually decreased the number of Las genes that could be detected although the total reads mapped to the Las genome were drastically increased. This is likely due to the loss of low-abundant Las gene transcripts during the treatment but significant enrichment of Las genes with relatively high abundance. Compared to the RNA enrichment treatments, RNA extracted from the bacterial cell isolation samples was effective in detecting a higher number of Las transcripts with up to 10-fold increase in read depth and coverage. For RNA extracted from ACP, the non-treated samples already had sufficient coverage (up to 100%) of the Las genome and the RNA enrichment treatments did not improve the performance.

**Table 2.**
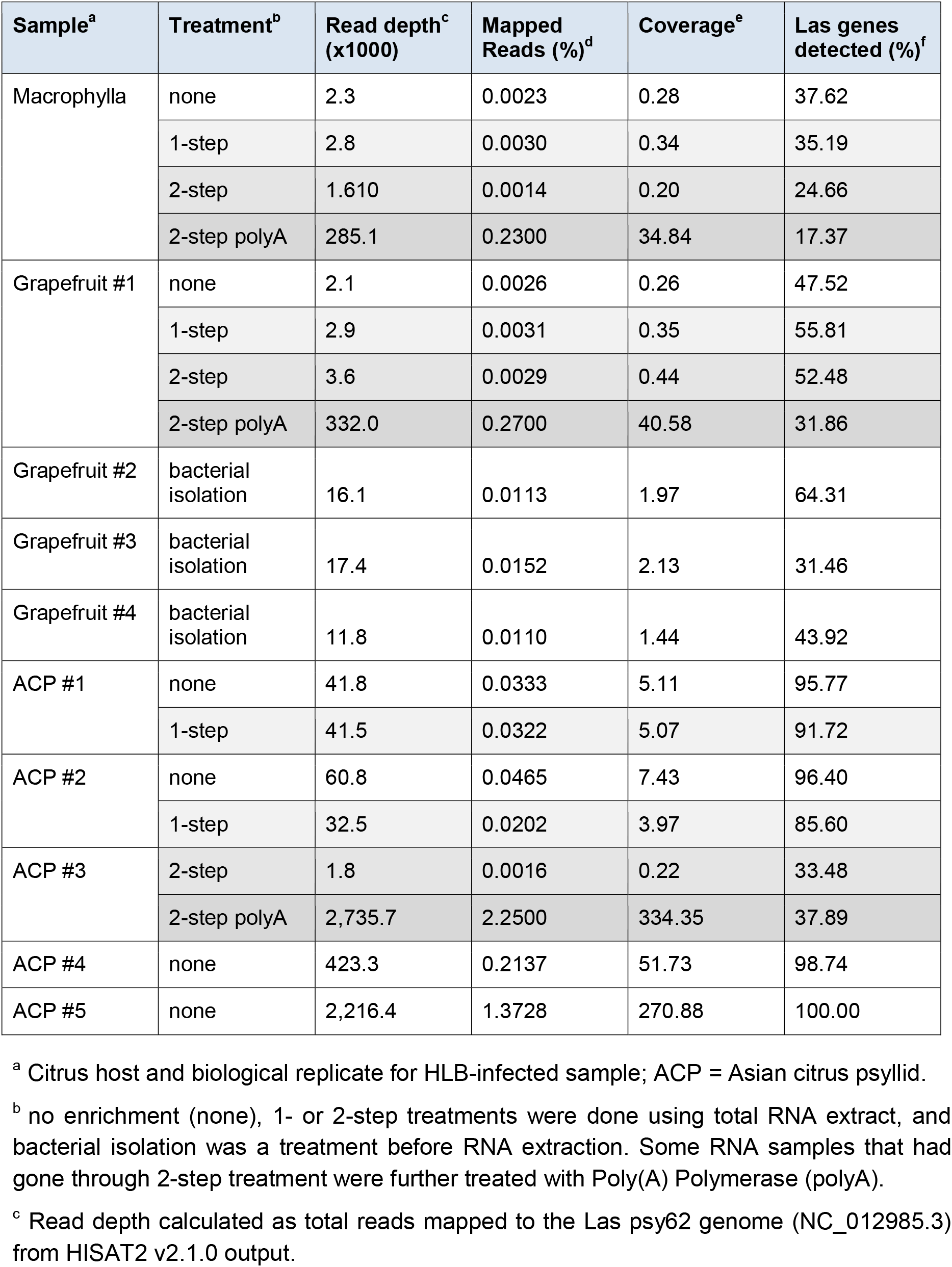
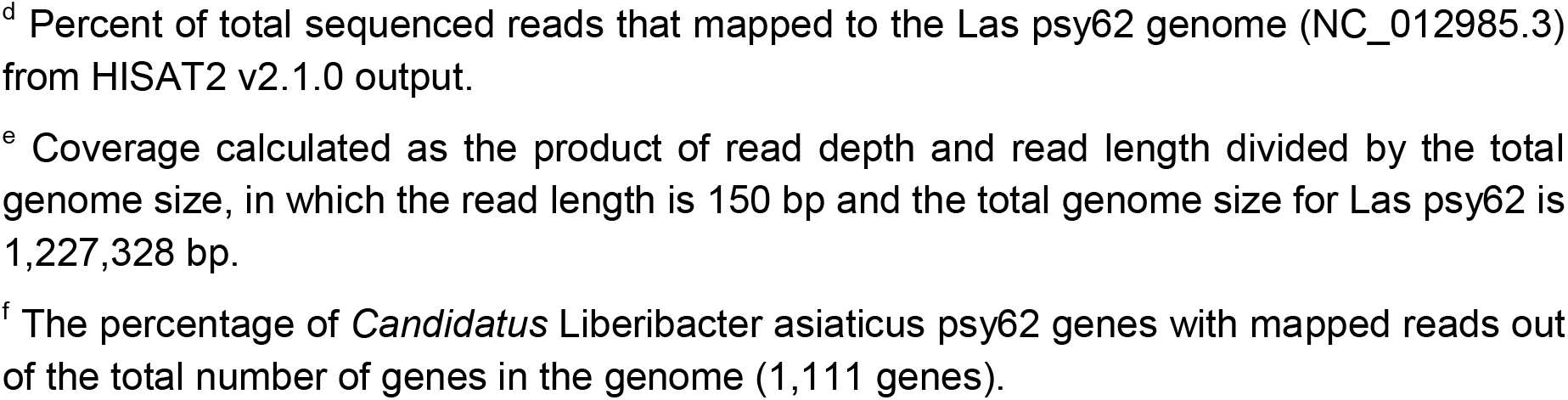
RNA-seq results using samples with various enrichment treatments.

| Sample<sup>a</sup> | Treatment<sup>b</sup> | Read depth<sup>c</sup><br>(x1000) | Mapped<br>Reads (%)<sup>d</sup> | Coverage<sup>e</sup> | Las genes<br>detected (%)<sup>f</sup> |
| --- | --- | --- | --- | --- | --- |
| Macrophylla | none | 2.3 | 0.0023 | 0.28 | 37.62 |
|  | 1-step | 2.8 | 0.0030 | 0.34 | 35.19 |
|  | 2-step | 1.610 | 0.0014 | 0.20 | 24.66 |
|  | 2-step polyA | 285.1 | 0.2300 | 34.84 | 17.37 |
| Grapefruit #1 | none | 2.1 | 0.0026 | 0.26 | 47.52 |
|  | 1-step | 2.9 | 0.0031 | 0.35 | 55.81 |
|  | 2-step | 3.6 | 0.0029 | 0.44 | 52.48 |
|  | 2-step polyA | 332.0 | 0.2700 | 40.58 | 31.86 |
| Grapefruit #2 | bacterial<br>isolation | 16.1 | 0.0113 | 1.97 | 64.31 |
| Grapefruit #3 | bacterial<br>isolation | 17.4 | 0.0152 | 2.13 | 31.46 |
| Grapefruit #4 | bacterial<br>isolation | 11.8 | 0.0110 | 1.44 | 43.92 |
| ACP #1 | none | 41.8 | 0.0333 | 5.11 | 95.77 |
|  | 1-step | 41.5 | 0.0322 | 5.07 | 91.72 |
| ACP #2 | none | 60.8 | 0.0465 | 7.43 | 96.40 |
|  | 1-step | 32.5 | 0.0202 | 3.97 | 85.60 |
| ACP #3 | 2-step | 1.8 | 0.0016 | 0.22 | 33.48 |
|  | 2-step polyA | 2,735.7 | 2.2500 | 334.35 | 37.89 |
| ACP #4 | none | 423.3 | 0.2137 | 51.73 | 98.74 |
| ACP #5 | none | 2,216.4 | 1.3728 | 270.88 | 100.00 |

<sup>a</sup> Citrus host and biological replicate for HLB-infected sample; ACP = Asian citrus psyllid.
<sup>b</sup> no enrichment (none), 1- or 2-step treatments were done using total RNA extract, and bacterial isolation was a treatment before RNA extraction. Some RNA samples that had gone through 2-step treatment were further treated with Poly(A) Polymerase (polyA).
<sup>c</sup> Read depth calculated as total reads mapped to the Las psy62 genome (NC\_012985.3) from HISAT2 v2.1.0 output.
<sup>d</sup> Percent of total sequenced reads that mapped to the Las psy62 genome (NC\_012985.3) from HISAT2 v2.1.0 output.
<sup>e</sup> Coverage calculated as the product of read depth and read length divided by the total genome size, in which the read length is 150 bp and the total genome size for Las psy62 is 1,227,328 bp.
<sup>f</sup> The percentage of *Candidatus Liberibacter asiaticus* psy62 genes with mapped reads out of the total number of genes in the genome (1,111 genes).

**Fig 2.**
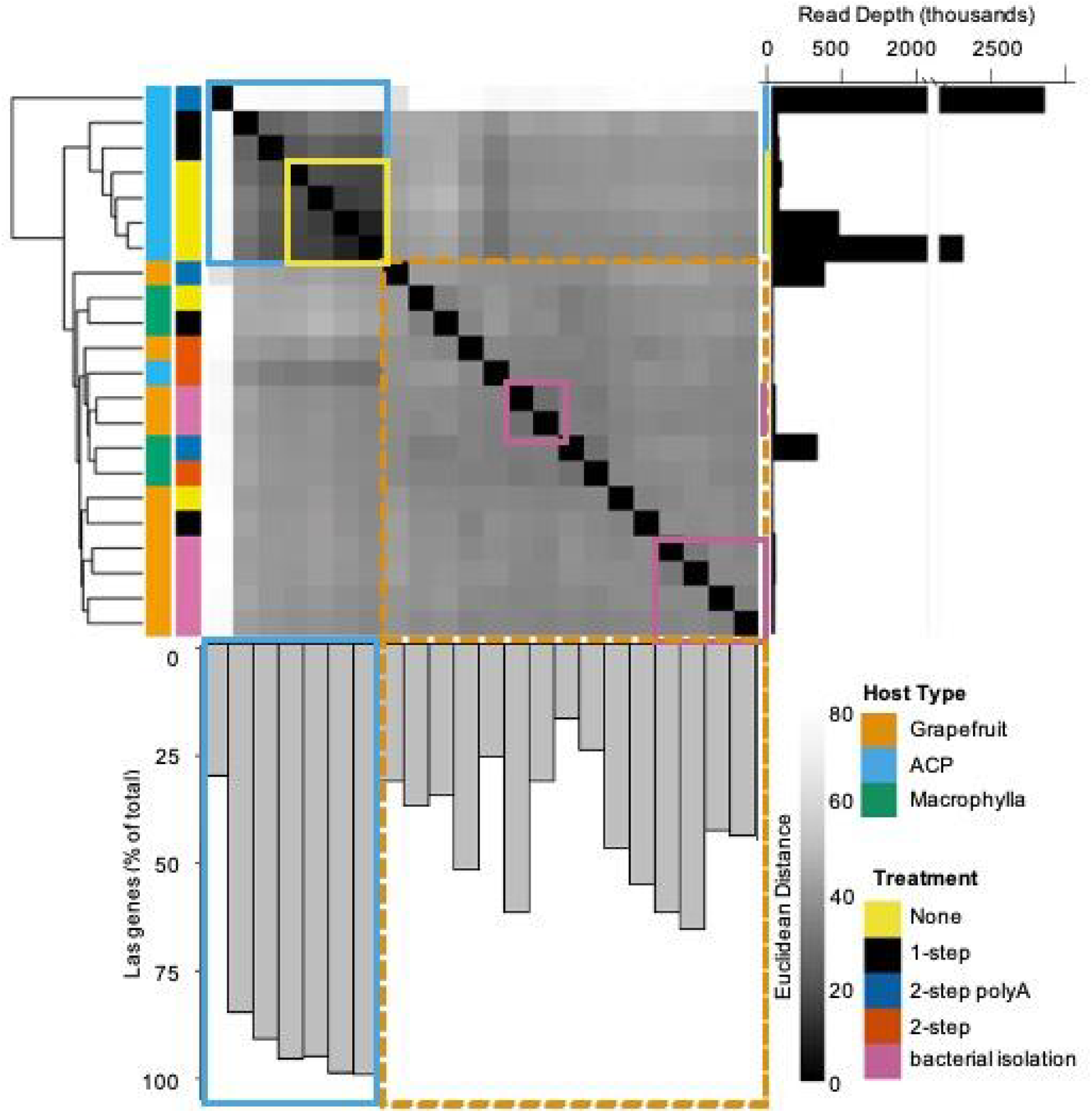
Summary of RNA-seq results after mapping to the Las psy62 genome. RNA-seq results of gene expression profiles were clustered based on Euclidean distance. The read depth or number of reads mapped to the Las genome as well as the percent of total genes with mapped reads are plotted for each sequenced sample. The citrus and ACP samples are designated by a dotted orange and solid blue box respectively. Grapefruit samples that underwent bacterial isolation and ACP samples that received no treatment are designated by a pink and yellow box respectively to indicate samples subjected to further analysis.

Euclidean distance analysis and principal component analysis of the log_2_-transformed gene counts for all samples and treatments were performed for quality assessment and to statistically distinguish them in terms of overall gene expression profile (Fig. 2, Fig. 3, Supplemental Data Set 1). The results revealed that the samples grouped together largely based on the host type from which the samples were isolated from. Samples collected from ACP and citrus have distinct transcriptome profiles. For example, all macrophylla samples clustered together in the PCA regardless of bacterial isolation or RNA enrichment, indicating that the bacterial or RNA enrichment treatments did not alter Las gene expression profiles. The 1-step and 2-step treated samples showed similar transcriptome profiles to the untreated controls, further confirming that they were ineffective in enriching Las RNA. A shift in transcriptome profiles in both citrus and ACP was observed after 2-step (PolyA) treatment, likely due to the substantial loss of low-abundant Las RNA.

**Fig 3.**
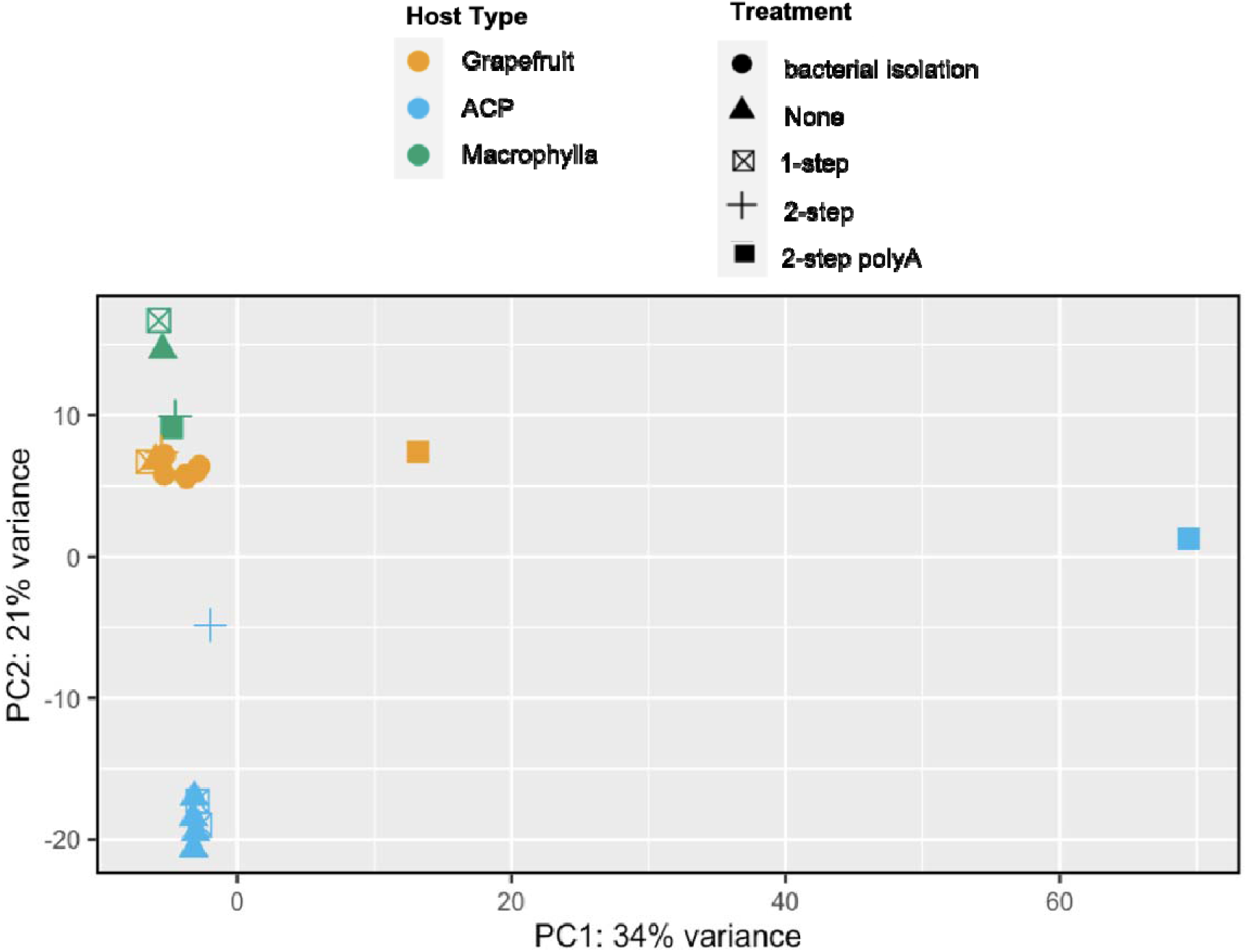
Principal component analysis of gene expression profile measured in log-transformed bacterial gene counts for HLB-infected hosts. Hosts included in the analysis include grapefruit (orange), macrophylla (green), and Asian citrus psyllid (ACP, blue). Samples were subjected to different treatments: no treatment (none, triangle), bacterial isolation (circle), and RNA enrichment: 1-step (crossed-square), 2-step (cross), and 2-step polyA (square). Gene counts from RNA-seq data were transformed using the rlog function in DESeq v.1.30.1.

Based on the sequencing results and the clustering analyses, we concluded that the RNA extracted from bacterial isolation samples of citrus and the untreated RNA extracted from ACP samples provided the best RNA-seq data sets for each host type. More specifically, these samples gave the most consistent results among biological replicates, demonstrated by the tightest clustering, the best read depth/coverage, and the largest number of Las genes detected. Therefore, these data sets were used for further gene expression analysis.

### Las genes with significant expression in citrus

Compared to ACP, Las has a particularly detrimental impact on citrus; therefore, we analyzed the Las genes with detectable expression in citrus and focused on the highly expressed genes, which indicate that they may play important roles in colonizing citrus. The distribution of an average of normalized read counts (counts per million or CPM) was determined among citrus samples (three biological and two technical replicates, Fig. 4A). We counted the number of Las genes that fell into 14 groups within the range of CPM from 0 to 15,675. Most genes belong to the first category with a CPM of 0-4. About 140 genes have the average CPM between 4 and 8 and 96 genes have an average of CPM ≥ 8 (Supplemental Table 2). This latter group represents the top 10% highest expressed genes within the Las genome. A GO enrichment analysis was performed on these 96 genes, from which 40 were found to be enriched in one or more GO terms. These genes were distributed among several enriched GO terms as shown in Figure 4B (Supplemental Data Set 3). Many of these genes are linked to housekeeping activities, including primary metabolism, macromolecule biosynthesis and gene expression. Genes with functions in stress responses, including chaperones and heat shock proteins, and mobile element-related activities, such as transposases, phage capsid, and other phage-related genes, were also enriched. Among other genes worth highlighting are ferritin, ion transporters, outer membrane porins, and genes for pilus formation and cell wall modification. In particular, a potassium transporter (*kup*) was among the highest expressed Las genes in citrus (Fig. 4A).

**Fig 4.**
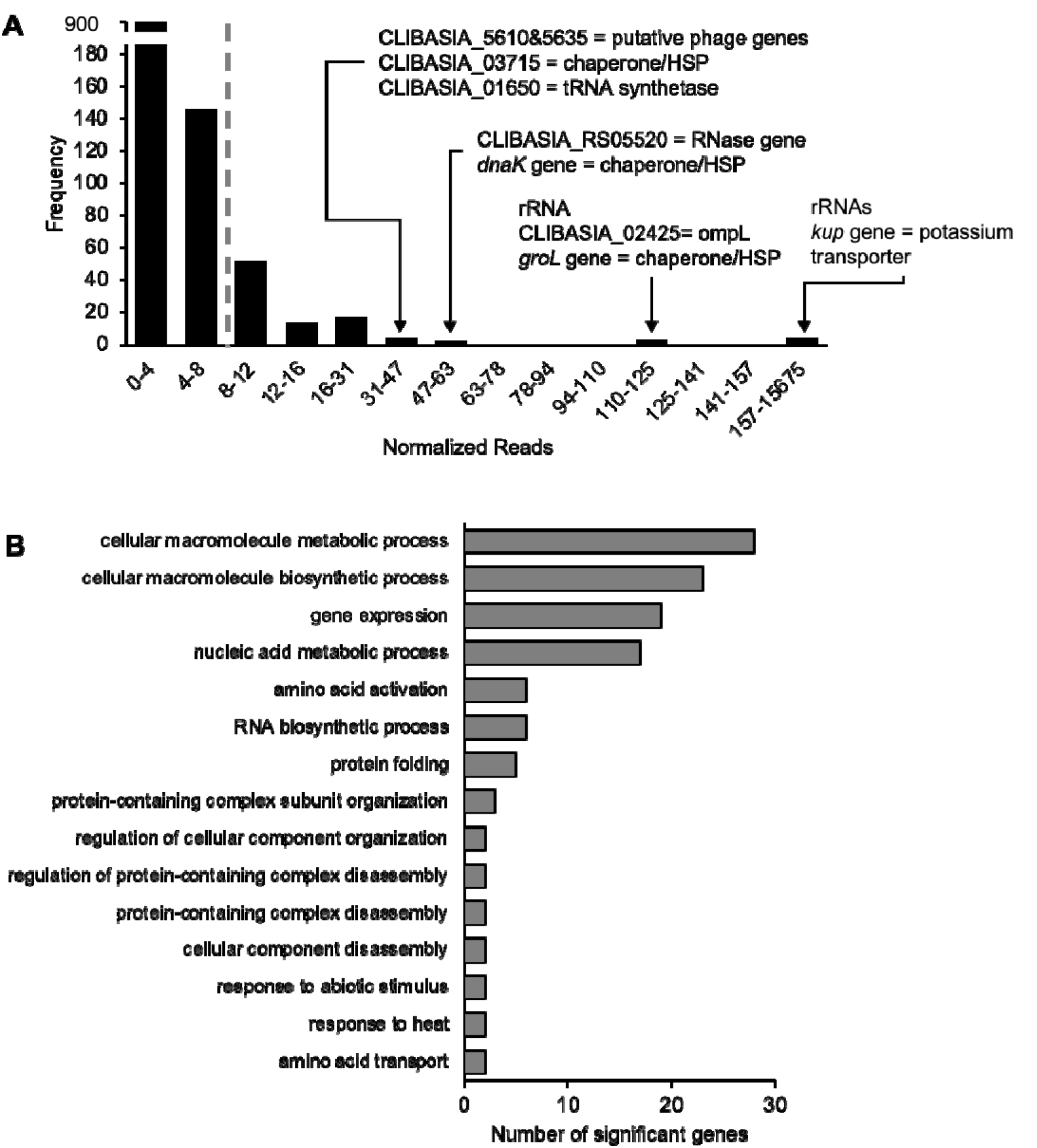
Highly expressed Las gene in citrus. **(A)** Distribution of normalized read counts mapped to the Las psy62 genome where frequency refers to the number of Las genes with average CPMs that fall within each bin. The top 96 genes (with average CPM ≥ 8) were selected for GO enrichment analysis and can be found in Supplemental Table 2. **(B)** List of non-redundant significantly enriched GO terms found in the top 96 Las genes expressed in citrus. The number of significant genes for each GO term are plotted. For a full GO list, see Supplemental Data Set 3.

As for other genes previously reported to associate with Las infection and HLB development in citrus, the gene CLIBASIA_05590, a prophage-encoded peroxidase (SC2_gp095) (Jain et al. 2015), has an average of CPM of 3.9, and the gene CLIBASIA_00255, encoding the salicylic acid hydroxylase (*sahA*) (Li et al. 2017), has an average of CPM of 1.35. In addition, a Sec-dependent effector (SDE) candidate CLIBASIA_03230 showed the highest expression level (CPM of 12.6) compared to other SDEs and is the only SDE that is among the top 10% expressed Las gene in citrus.

### Differential expression of Las genes in citrus and psyllids

We compared the transcriptome profiles of Las in citrus (midrib of grapefruit) and ACP in order to investigate which genes might be important for the HLB disease cycle. Reads per gene were tabulated and statistical analyses were performed. Of the 1,111 annotated genes in the Las genome, 931 and 1,111 had detectable expression in citrus and ACP samples respectively, corresponding to 84% and 100% coverage. Hierarchical clustering of the complete gene set encoded by the Las genome based on their relative expression in each host grouped the genes into ten clusters, each exhibiting a different expression pattern. After excluding 409 genes, which had low read counts in citrus that are insufficient to support robust statistical analysis, we identified 106 genes with significant differences (|log_2_FC| > 0 and padj < 0.05) in relative expression between hosts (Fig. 5, Supplemental Data Set 2). Most of these differentially expressed genes (DEGs) are distributed across clusters I-VIII with the majority of the DEGs found within clusters VI and VIII.

**Fig 5.**
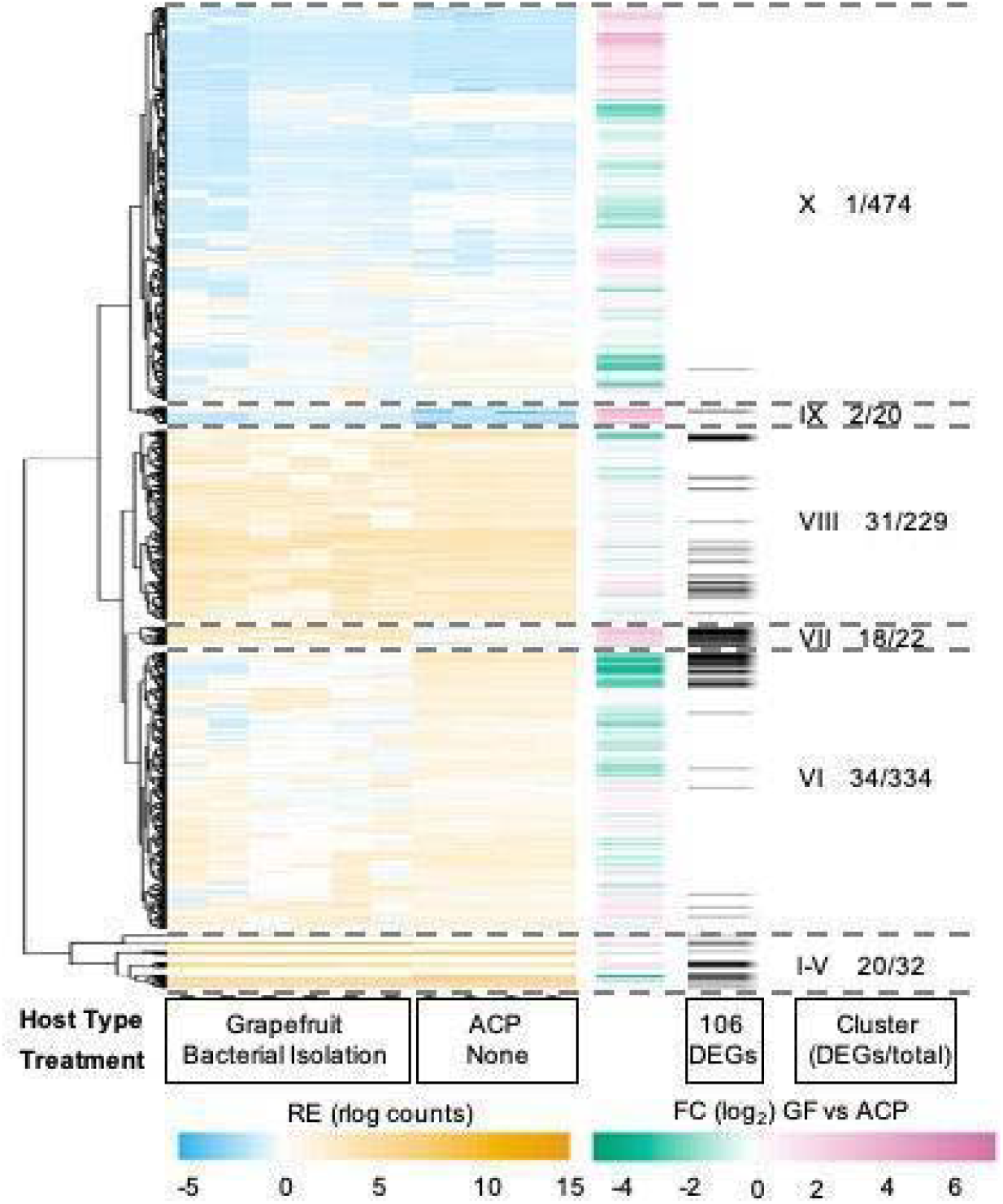
Differentially expressed Las genes in citrus vs ACP. (Left; blue/orange heatmap) Hierarchical clustering of the relative expression (RE) of Las psy62 genes in grapefruit (GF) midrib samples after bacterial isolation pre-enrichments and Asian Citrus Psyllid (ACP) samples with no enrichment. (Middle; green/purple heatmap) Fold changes in Las gene expression in GF samples compared with ACP samples (Right; black/white heatmap). Differentially expressed genes (DEGs) represented by the black bars based on comparisons between GF and ACP samples where DEGs were identified as genes with |log_2_ fold change| > 0 and an adjusted *p* value < 0.05. Heatmaps were divided into ten clusters where the total number of genes and total number of DEGs are reported for each cluster.

Next, the genes in each cluster and the DEGs were subjected to GO analysis (Fig. 6, Supplemental Data Set 3). Significantly enriched GO terms were identified for each set of genes. In cluster VII, genes related to defense and interspecies interaction were significantly induced in Las during infection of citrus compared to Las found in ACP. These genes include lysozyme-encoding genes CLIBASIA_04790 and CLIBASIA_04800, which could play a role in the restructuring of the bacterial cell wall in response to a defense response *in planta.* Similarly, in cluster VIII, genes related to transcription, biosynthetic processes, cellular macromolecule metabolism, and translation were also upregulated in citrus.

**Fig 6.**
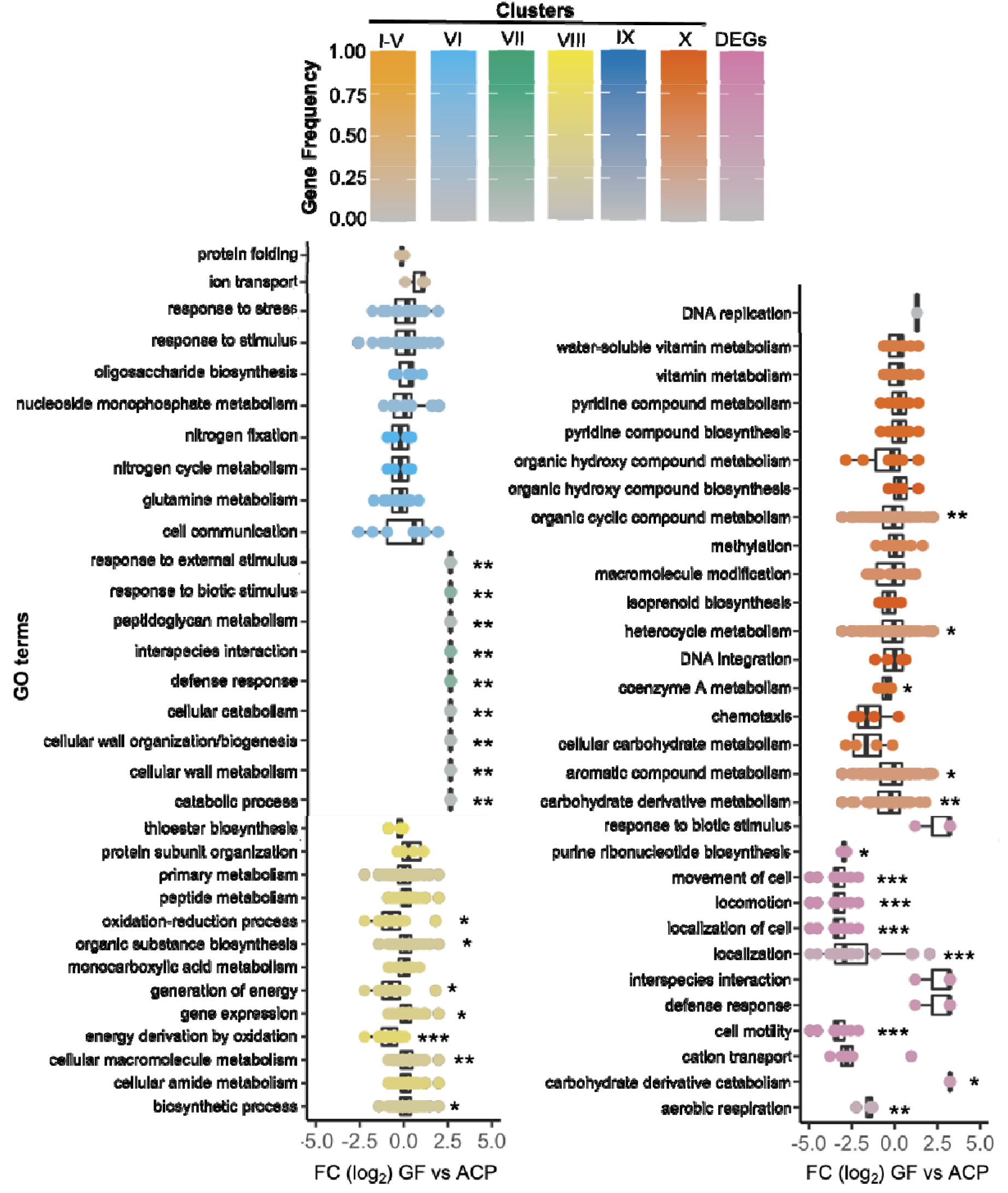
Non-redundant significantly enriched GO terms found in Las gene clusters and differentially expressed genes (DEGs). Significant genes for each GO term are plotted based on their fold changes (log_2_FC) in Las gene expression in citrus samples compared with ACP samples. The gene frequency represented as # of significant genes / total number of genes for a given GO term gene group is given by the dotplot color. For a full GO list, see Supplemental Data Set 3. * indicates log_2_FC values significantly different than 0 (2-tailed t-test at * *p* < 0.05, ** *p* < 0.01, *** *p* < 0.001).

In analyzing the 106 DEGs for significantly enriched GO terms, we found that genes involved in motility and aerobic respiration were repressed in Las isolated from citrus compared to Las colonizing ACP. More specifically, the significantly expressed genes from these GO terms belong to the TCA cycle enzymes such as succinate dehydrogenase as well as NADH dehydrogenase complexes. Succinate dehydrogenase is a membrane-bound enzyme of the TCA cycle which is involved in the aerobic electron transport pathway to generate energy via oxidative phosphorylation pathways. NADH dehydrogenase is also a membrane-bound enzyme involved in oxidative phosphorylation in the electron transport chain to generate energy in the form of ATP. KEGG pathway analysis supports these findings in that genes involved in the flagella assembly pathway and the oxidative phosphorylation pathways were significantly downregulated in citrus (q < 0.05) (Supplemental Data Set 4). For example, the *flaA* gene (CLIBASIA_02090) that encodes flagellin and ten additional flagella-related genes were highly expressed in ACP. *flaA* has a low expression in citrus (average CPM = 1.83) but showed a log_2_FC of −4.52 in citrus vs ACP, consistent with the previous report that flagella were only observed in ACP but not in plant hosts (Andrade et al. 2020).

### Correlation analysis of the RNA-seq results with previously published RT-qPCR data

RT-qPCR analysis has been used to investigate relative expression of individual Las genes in citrus vs ACP. To determine how our transcriptome data align with the previously published RT-qPCR results, we conducted correlation analyses using log_2_FC values between *in planta* and in psyllid expression (Fig. 7). In particular, we compared our RNA-seq data with two published studies, which analyzed 198 Las genes (Yan et al. 2013) and 17 SDE genes (Thapa et al. 2020) respectively. Ten of the 198 genes analyzed in Yan et al. (2013) were not included in this analysis due to re-annotation of the Las genome. Correlation analysis between the entire gene set from Yan et al. (2013) and our RNA-seq analysis was significant (*p* < 0.05) based on a Spearman’s test which indicates that there is a relationship between the two datasets; however, with a correlation coefficient value of 0.15, this correlation is weak. The genes analyzed in Yan et al. (2013) were categorized based on specific pathways, so we analyzed the correlation coefficient value for each set of genes separately. Of the seven categories, genes involved in the secretion system had a significant positive correlation with our RNA-seq data (*p* < 0.05, R = 0.36). Similarly, correlation analysis with the RT-qPCR results of predicted Las SDEs from Thapa et al. (2020) also revealed a significant positive correlation (*p* < 0.05, R = 0.52). The lack of correlation in other gene sets is probably due to the difference in the experimental approach and samples being analyzed.

**Fig 7.**
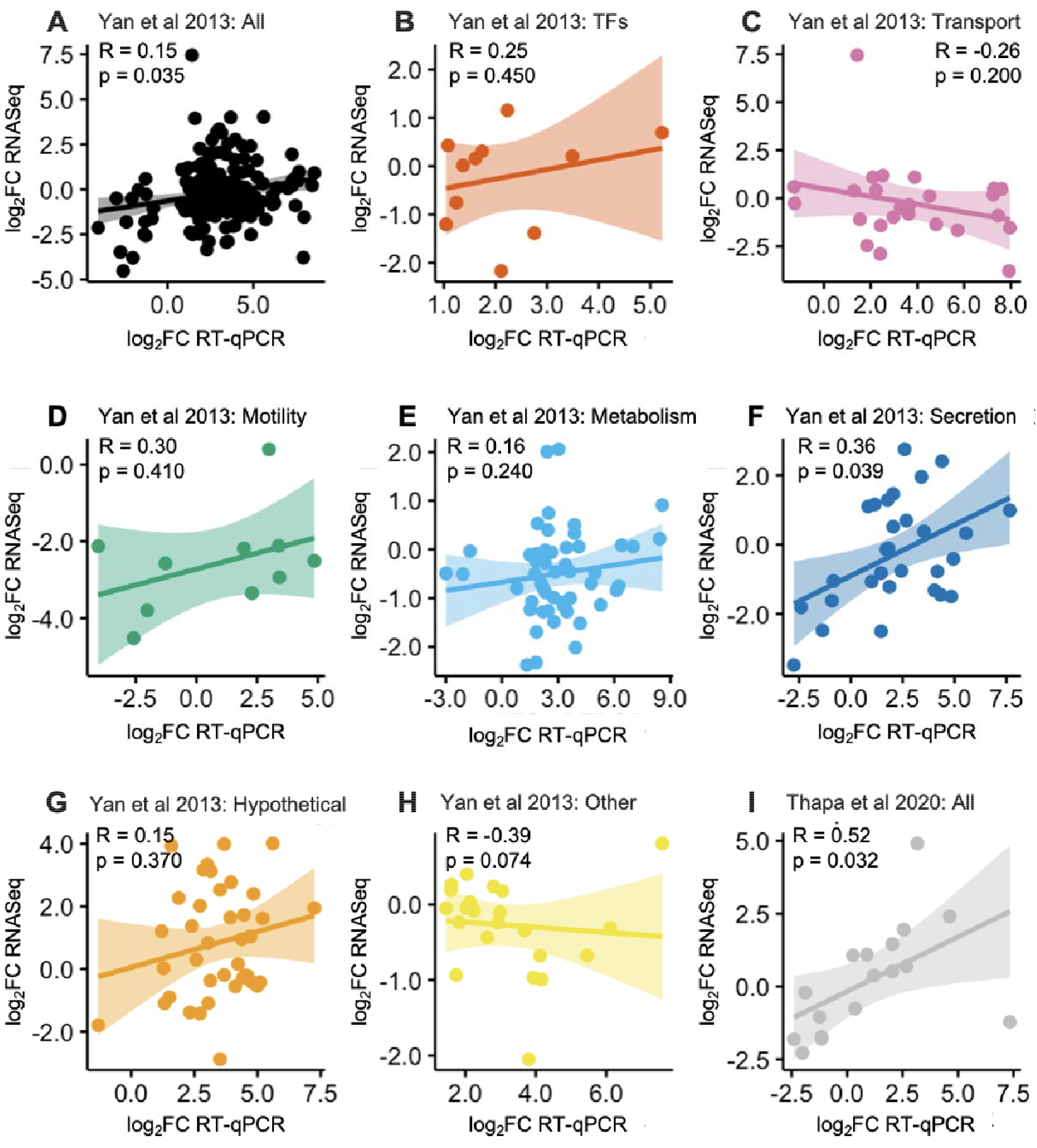
Comparison of RNA-seq results with previously published RT-qPCR data. Relationship between log_2_ fold changes (log_2_FC) values in Las gene expression in citrus samples compared with insect samples from our RNA-seq analysis (y-axis) and previously published RT-qPCR analyses (x-axis). Genes were grouped based on categories defined by Yan et al. (2013) including **(A)** All 188 genes, **(B)** Transcription Factors (TFs), **(C)** Transport systems, **(D)** Motility, **(E)** Metabolism pathways, **(F)** Secretion system, **(G)** Hypothetical proteins, and **(H)** Other. **(I)** Predicted core effectors defined in Thapa et al. (2020). Points represent relative expression of genes, lines represent the regression lines, shaded areas represent the 95% confidence interval. Spearman’s correlation coefficient, R, along with calculated p-value (p), given for all Yan et al. (2013) comparisons. Pearson’s correlation coefficient, R, along with calculated p-value (p), given for Thapa et al. (2020) comparison.

### Expression profiles of predicted SDEs

As effectors play a key role in disease development, we examined the gene expression levels of the 27 predicted core SDEs defined by Thapa et al. (2020) in citrus and ACP (Supplemental Table 3). Since CLIBASIA_01000, _03120 and _04970 were not annotated in the newer CLIBASIA_RS system and thus not included in the RNA-seq data, the remaining 24 core SDEs were analyzed. Fifteen of these SDEs were expressed in both ACP and citrus, eight were only expressed in ACP, and one (CLIBASIA_00460) was only expressed in citrus (Fig. 8). Of the 15 SDEs expressed in both citrus and ACP, two were identified as significant DEGs, including CLIBASIA_05315 (or SDE1) showed significantly higher expression in citrus with log2FC = 4.91; and CLIBASIA_04410 exhibited significantly higher expression in ACP with log2FC = −1.95.

**Fig 8.**
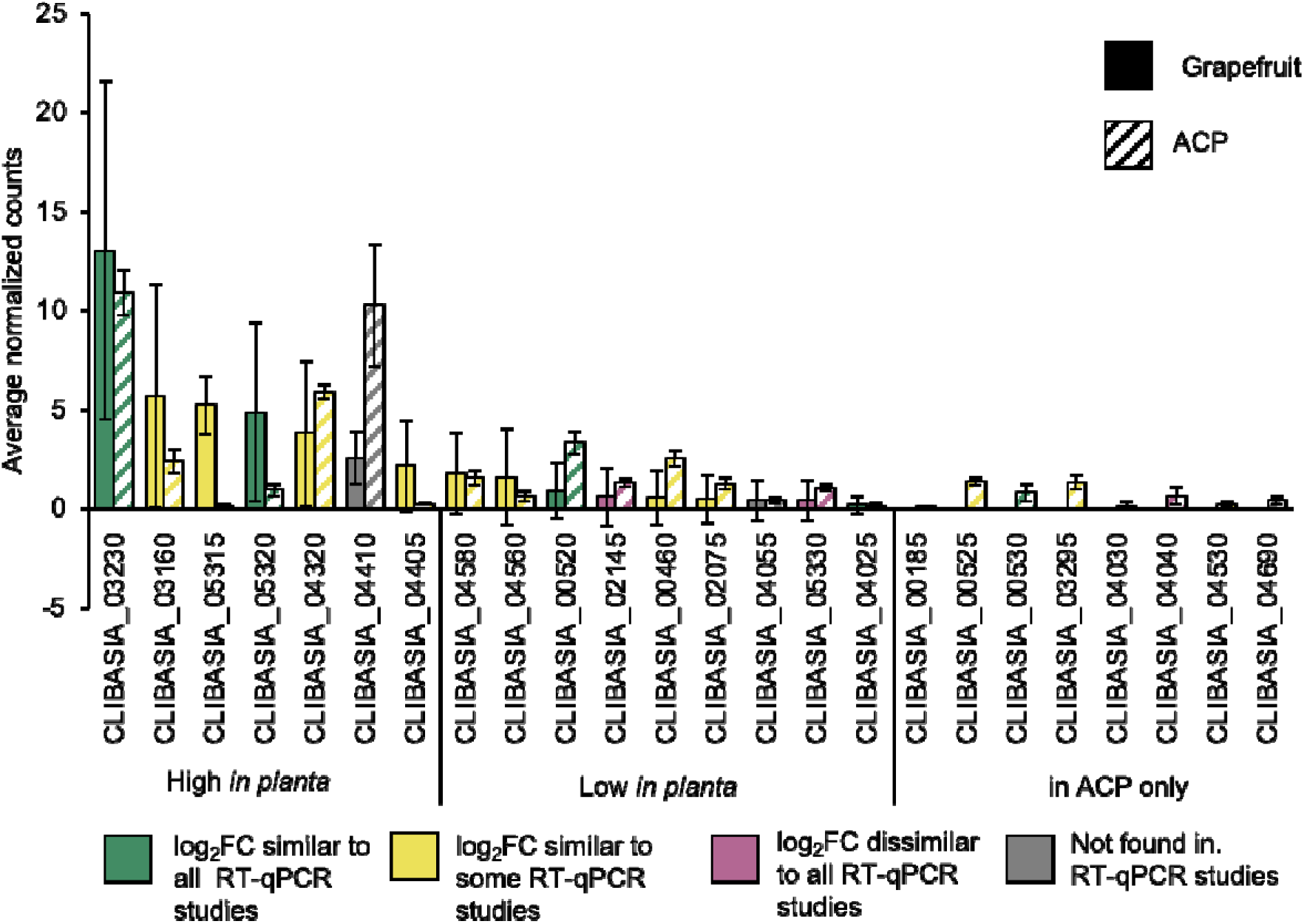
Expression of predicted core Sec-Dependent Effectors (SDEs) in citrus and psyllid. The average normalized read counts for each predicted core SDE in citrus (Grapefruit, solid) and Asian citrus psyllid (ACP, striped) samples listed in order based on relative *in planta* expression levels. Genes divided based on relatively high (> 0 reads, *p* < 0.05), low (> 0 reads, *p* > 0.05) or no (= 0 reads) using RNA-seq data in citrus where significance is based on 2-tailed t-test. Color responds to the correlation with log_2_FC values reported in RT-qPCR studies Yan et al. (2013) and Thapa et al. (2020).

When comparing the relative expression of the 24 SDEs in citrus, seven genes were classified as significantly expressed, of which CLIBASIA_03230 had the highest expression in both citrus and ACP (Fig 8). Because many core SDEs showed relatively low expression, we examined their expression in citrus using RT-qPCR by comparing Ct values between infected and healthy tissues (Supplemental Table 3). Out of the eight SDEs that were undetectable in citrus by RNA-seq, five were detectable by RT-qPCR (Supplemental Table 3), indicating that they were probably expressed but at a very low level. Three SDEs were undetectable by both RNA-seq and RT-qPCR, suggesting that they were not expressed, at least in the samples used in this study.

## Discussion

This work utilized a range of new and rigorous methods aimed at establishing the transcriptome landscape of Las in its hosts. We were able to achieve a much improved coverage of the Las genome by RNA-seq using a bacterial isolation protocol modified from Nobori et al. (2018). RNA enrichment treatments aiming to increase the proportion of Las RNA in total RNA extracts from infected citrus and ACP tissues were also tested. A similar approach was successfully used in other host-microbe systems (Kumar et al. 2016; Lovelace et al. 2018) but it was insufficient to enrich Las RNA. This is likely due the extremely low abundance of Las RNA in the samples and the inevitable loss of the low-abundance RNA during the process. Taken together, for infected samples with low bacterial titer like in the case of Las-infected citrus tissues, the best practice would be to use bacterial cell enrichment rather than RNA enrichment for bacterial transcriptome analysis.

Bacterial accumulation in the host types tested in this work was critical to recover sufficient Las genome coverage by RNA-seq. In the case of citrus, we used two species: *Citrus paradisi* Macfadyen (Rio Red grapefruit) and *Citrus macrophylla*. Samples were collected from naturally infected grapefruits from fields where the disease has effectively established, presumably with Las strains infections (Sétamou et al. 2020). Macrophylla is a commonly used rootstock, where HLB-infected trees showed higher bacterial titers but less severe symptoms (Folimonova et al. 2009; McCollum et al. 2016). For both species, we selected highly infected samples with clear HLB symptoms, likely representing a late infection stage.

Based on our RNA-seq analysis, 96 Las genes showed the highest expression levels (representing top 10%) in citrus. The majority of these genes belong to pathways that are important for basal cellular functions including gene regulation, transcription, translation, metabolism, and energy production. Recently, Fang et al. (2021) reported comparative Las gene expression in citrus midrib with fruit pith using dual-transcriptome data. Many genes found to be highly expressed in citrus in their study were also found in this work. For example, ferritin, lysozymes, porins, and chaperones were found in both studies, indicating a role of these genes in the life cycle of Las with possible contributions to disease development. As one of the most highly expressed Las genes in citrus, the ferritin gene, CLIBASIA_03035, encodes an oxidase that detoxifies excess ferric ions. Iron metabolism and iron-immunity crosstalk were found to be essential in plant immune response (Herlihy et al. 2020; Inoue et al. 2020). It is therefore possible that CLIBASIA_03035 may manipulate this process and promote disease. Some porins and transporters were highly expressed in citrus. Among them is the outer membrane immunogenic protein OmpL (CLIBASIA_02425). The outer membrane porin OmpA (CLIBASIA_03315) has been used as a detection marker for HLB diagnosis (Ding et al. 2015). With a higher expression level than OmpA, OmpL could serve as another biomarker to enhance HLB detection. CLIBASIA_01685, which encodes a Bax inhibitor-1/YccA protein, is also highly expressed. The BAX inhibitor is known to have the conserved function as a cell death suppressor in mammals and plants (Hückelhoven 2004). As an obligate pathogen, inhibition of cell death could be critical for the colonization of Las in the hosts. Furthermore, overexpression of YccA in *E. coli* inhibits FtsH-mediated degradation of SecY, a key component of the Sec secretion system (Stelton et al. 2009). As such, it is perceivable that CLIBASIA_01685 may promote the Sec secretion system and enhance the activity of SDE effectors to promote infection. Finally, the gene *flp1* (CLIBASIA_03115) encodes one of the core Flp pilins (fimbrial low-molecular-weight proteins) found in the Las genome (Andrade and Wang, 2019). Pili have a broad range of functions in bacterial attachment, movement and substrate transport (Hospenthal et al. 2017). In addition, the type IVc tight adherence (Tad) pilus apparatus may contribute to Las colonization of psyllids (Andrade and Wang 2019). It is therefore interesting to hypothesize that Flp1 may facilitate the colonization of Las in citrus.

Las engages in complicated interactions with both citrus and ACP (Galdeano et al. 2020; Huang et al. 2020). In addition to manipulating citrus, Las also causes metabolic alterations in ACP, increasing their mortality rate (Killiny et al. 2017). We found 106 Las genes that are differentially expressed in citrus vs ACP. These genes are distributed throughout the Las genome. Half of them (53) were found as upregulated in citrus and the other half (53) in insects. Genes related to defense and interspecies interaction were upregulated in citrus. These genes may promote bacterial survival during the infection, either by counteracting plant defense responses or protecting against bacteriophages (Forsberg and Malik 2018). Among this group are genes encoding lysozymes whose immunomodulatory functions in the context of infection might explain their significant upregulation in citrus (Ragland and Criss 2017). Genes involved in interspecies interaction might be relevant for establishing communication with other microbes within a microbial community. These interactions can be beneficial, syntrophic or competitive, and indeed determine the fitness and evolution of Las in its environment (Riera et al. 2017; Gorter et al. 2020; Blacutt et al. 2020).

Another important group of DEGs was found to be involved in gene expression, biosynthetic processes, cellular macromolecule metabolism, and organic substance biosynthesis such as ribosomal proteins and tRNAs. All of them were significantly upregulated in citrus, suggesting that transcription and translation are more active when Las colonizes citrus, consistent with the assumption that Las infection involves genetic reprogramming *in planta*. However, the activation of these processes in citrus did not correlate with an increase in metabolic activity. On the contrary, it is interesting that genes coding for enzymes participating in energy generation were upregulated in psyllids. This observation can be related to the previous findings which suggests that Las stimulates ACP to produce more ATP and other energetic nucleotides (Killiny et al. 2017). The TCA cycle was another energy-generating pathway found as upregulated in ACP. It is interesting to note that Kruse et al. (2017) reported that the expression of ACP genes involved in the TCA cycle was also significantly changed with Las exposure. Another gene cluster that was significantly upregulated in ACP is the flagella system. This is consistent with the previous reports on the downregulation of Las flagellin in citrus (Yan et al. 2013) and formation of flagella only in psyllids (Andrade et al. 2020).

The RNA-seq data provide novel insight into effector biology in Las. CLIBASIA_05315 or SDE1 was shown to interact with papain-like cysteine proteases in citrus (Clark et al. 2018), promote symptom development (Pitino et al. 2018; Clark et al. 2020), and used as a biomarker for HLB detection (Pagliaccia et al. 2017). Consistent to its important virulence function in citrus, SDE1 is one of the most highly expressed SDEs and almost exclusively expressed in citrus. The SDE candidate CLIBASIA_04410, which encodes a hypothetical protein with predicted domains for a biotin-protein ligase, was expressed in both citrus and ACP but showed significantly higher expression in ACP. Another SDE candidate CLIBASIA_03230 is equally highly expressed in citrus and ACP. Encoding a hypothetical protein, CLIBASIA_03230 is the highest expressed SDE in both citrus and ACP. How SDEs may manipulate the insect vector is unknown. CLIBASIA_04410 and CLIBASIA_03230 could be good research subjects to start understanding this overlooked component of HLB biology. It would also be interesting to determine whether CLIBASIA_03230 targets similar cellular processes in plants and insects.

When comparing our RNA-seq data set with previously published RT-qPCR data sets, only secretion-related genes, especially SDEs, have significant positive correlations with our RNA-seq results. The lack of correlation in other gene sets may be due to differences in experimental designs: our *in planta* samples were collected from naturally HLB-infected, symptomatic, adult grapefruit trees found in Texas and macrophylla trees from Florida whereas the samples from Yan et al. (2013) were collected from symptomatic sweet orange seedlings which were infected via grafting. Differences in plant age and environmental conditions may also contribute to the little correlation between our RNA-seq data and the RT-qPCR results. Additionally, transcriptomics analyses by RNA-seq allows comparison between different genes under the same condition whereas PCR-based approaches can only compare the expression level of the same gene under different conditions.

In summary, the current study provides a significant contribution to the knowledge of Las gene expression profiles in naturally-infected, mature citrus trees at late infection stages and ACP. The established bacterial cell isolation protocol is a useful tool for similar investigations on the transcriptome of Las and other Liberibacter species, as well as other bacterial pathogens with low titers in infected tissue.

## Author Contributions

WM conceived and designed the experiments. CW, FG, VA and AL provided materials. ADF and MQ performed the experiments. TJ, DS and AHL did the bioinformatics and statistical analyses. ADF and AHL prepared figures/tables and wrote the manuscript with WM. All authors contributed to the article and approved the submitted version.

## Acknowledges

We thank the members of the Ma lab for their helpful discussions and feedback and USDA-NIFA for supporting this research.

## Notes

**Funding**: This work is supported by USDA National Institute of Food and Agriculture award No. 2016-70016-24833 to W.M and V.A, No. 2019-70016-29796 to W.M, and No. 2020-70029-33197 to W.M and A.L.

### Competing Interest Statement

The authors have declared no competing interest.

