## supplemental tables and figures for "Transcriptome profiling of *Candidatus* Liberibacter asiaticus in citrus and psyllids"

**Supplemental materials**

**Supplemental Fig 1 Post-enrichment steps deplete plant and bacterial rRNAs.** Bioanalyzer electropherograms of total RNA and bacterial mRNA-enriched samples from CLas-infected citrus grapefruit midribs. Signal intensity (FU) vs RNA size (nt). Pink arrows indicate the bacterial ribosomal RNAs ,16S and 23S and blue arrows the eukaryotic ribosomal RNAs 18S and 28S. The first plot (on top) shows the total RNA control with no enrichment (none) and the second and third plots show RNA after the bacterial post-enrichment (1- and 2-step respectively).

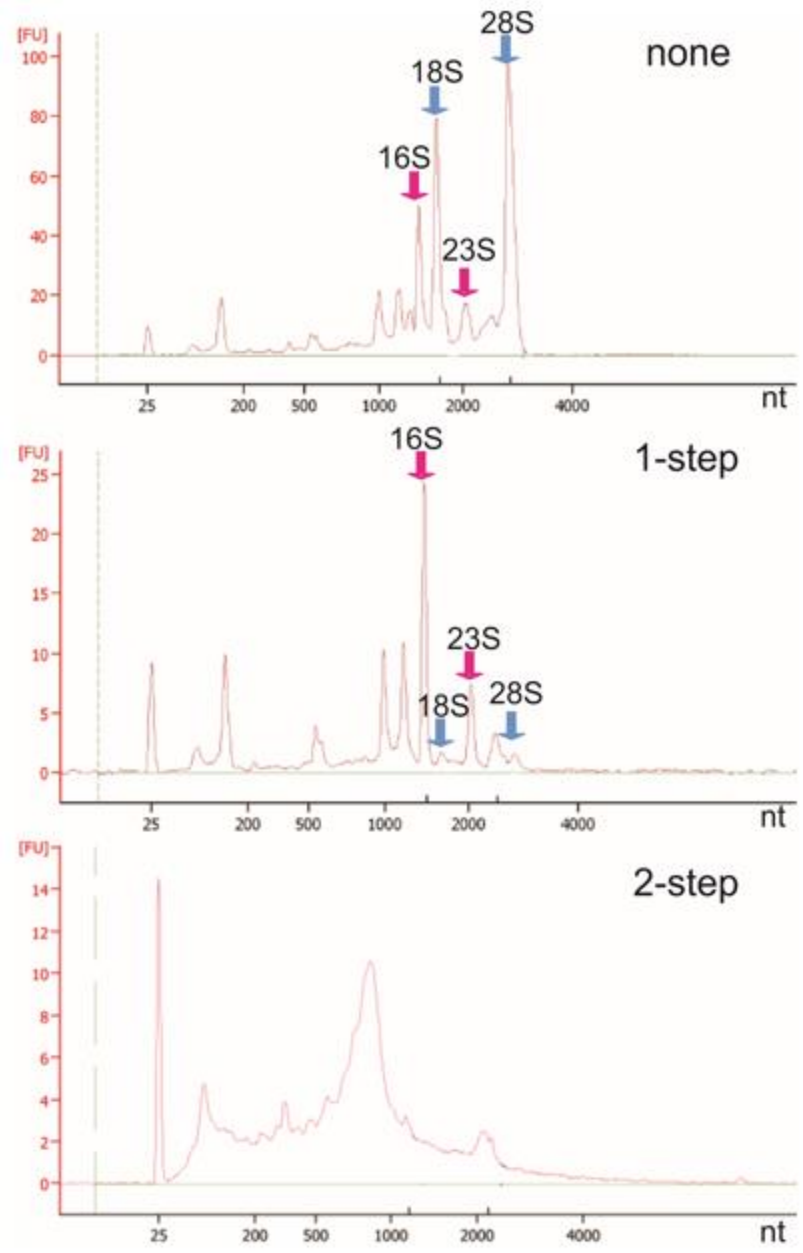


**Supplemental Table 1.** Primers sequences used in this work

| primer name/gene | F sequence | R sequence | Organism |
|---|---|---|---|
| CLIBASIA_00185 | TCCACATTAGCTGGCTGTGA | TCACCAAGGTCACGTCGATA | *Candidatus* Liberibacter asiaticus (Las) |
| CLIBASIA_00460 | GCCTCGTATTGCAACAAAATC | GTACACGGCGGAAGAAGATG | Las |
| CLIBASIA_00520 | CCTCTTAGGGAGCTGTGACG | GCAAGTTCCTTAGCGCTGTT | Las |

| | | | |
|---|---|---|---|
| CLIBASIA_00525 | ATCCGTCTACGCAAGACGAC | TGCAGATATTTCGGCTTCTG | Las |
| CLIBASIA_00530 | CTGTGCCGATGACAGGATTA | TGGACCATCCATACGTCTCA | Las |
| CLIBASIA_01000 | GGTGGTTCGCCATGTCTTAT | AAAAATATAAATCGAAGCGTAAAAA | Las |
| CLIBASIA_02075 | AAGTTGGAGAGCGTGCATTT | ATCGCTCGTTGCTCCTCTAA | Las |
| CLIBASIA_02145 | CCGAATTTGCTGGTATGGAT | GAGCTTCGCGATCATCTCTT | Las |
| CLIBASIA_03120 | ACTCAGATCGCATTGGCTTT | CGATGCAAAGGAGGTAAAGG | Las |
| CLIBASIA_03160 | GGTTTTGAACGGAGATCGTG | GACGCCCGTCATAACGTATT | Las |
| CLIBASIA_03230 | TGACGGGAATCAGTATCACTTTC | GCTAATGAACTTCAGAATAGCGA | Las |
| CLIBASIA_03295 | GAGTGAGTGTCGTTGGAGCA | AGACAACGCCTCTGCTGATT | Las |
| CLIBASIA_04025 | GGCATGATACTTTCTTCTTGTGG | TCACCATTCCATCCTCCTTC | Las |
| CLIBASIA_04030 | TCCAGTATTTGCAATGGGCACAGC | AAGAGCGACGGGAGCAGGAGGGATA | Las |
| CLIBASIA_04040 | AAGCAGCAGAACAAGCAGCAGAAG | ATTGGCTGCTACCGGTACCTCATT | Las |
| CLIBASIA_04055 | TTGCTCCGATCTTTGGAACT | GAAATAGCATTTCGAGCTCCTG | Las |
| CLIBASIA_04320 | AGGAGTCGGTGTCGTAGCAT | GAATCGCAAAATCGCTCTGT | Las |
| CLIBASIA_04405 | GAACGCTTTTTGGGATTTGA | TAATTTGATGGGGGCACAGT | Las |
| CLIBASIA_04410 | CACTGTCTGCGGAAAATGAA | GTAGCGGTGTCCGTTGTTTT | Las |
| CLIBASIA_04530 | ATGAGTGGGTGTTCCGAGAC | TTTCCCCTGTTTTTCCTCCT | Las |
| CLIBASIA_04560 | GTGACCTCGGTGATTCCATT | AGCCTGCTAGCACGACGTAT | Las |
| CLIBASIA_04580 | GTGGGTTATGCGAGCAATCT | CGGCAAAGGAGGTGGATATT | Las |
| CLIBASIA_04690 | ATGTGGATTGCTCCTTTTGG | AGAATGCCAGGGAATTGAGT | Las |
| CLIBASIA_04970 | ATGAACACAATGAAGAGAGAAAAA | TGCATAAACCTTTTTACCAATAGC | Las |
| CLIBASIA_05315 | CACCTAGAATCCATATGCGTCC | CCCCTAGAGCTGTTCATTTCAG | Las |
| CLIBASIA_05320 | TCTTAGCTGCCAATGAGCAC | GCCTCCAAAGAGCATAAGCA | Las |
| CLIBASIA_05330 | TATAACGTGCGGTGCACAAG | AAAGAATCAAGATTTTCTTTTGCTG | Las |
| 16S * | GGATAACGCATGGAAACGTGTGCT | AATCCAACGCAGGCTCATCTCTCT | Las |
| F-box** | TTGGAAACTCTTTCGCCACT | CAGCAACAAAATACCCGTCT | *Poncirus trifoliata* |

* (Yan et al. 2013)

** (Mafra et al. 2012)

**Supplemental Table 2.** Top highly expressed Las genes in citrus.

| Locus Tag[a] | Average NC[b] | Gene name | Gene Product | KEGG Pathway | PFAM domains[c] |
|---|---|---|---|---|---|
| CLIBASIA_r05780 | 15675 | | 23S ribosomal RNA | Ribosome | NA |
| CLIBASIA_r05782 | 14657 | | 23S ribosomal RNA | Ribosome | NA |
| CLIBASIA_03625 | 1206 | kup | putative potassium transporter Kup | Transporter | K_trans |

| | | | | | |
|---|---|---|---|---|---|
| CLIBASIA_r05785 | 273 | | 16S ribosomal RNA | Ribosome | NA |
| CLIBASIA_02425 | 122 | | porin family protein | Transporter | ompA_membrane |
| CLIBASIA_r05783 | 120 | | 16S ribosomal RNA | Ribosome | NA |
| **CLIBASIA_03720** | 112 | groL | chaperonin GroEL | RNA degradation | Cpn60_TCP1 |
| CLIBASIA_RS05520 | 56 | rnpB | RNase P RNA component class A | NA | NA |
| **CLIBASIA_02620** | 47 | dnaK | molecular chaperone DnaK | RNA degradation | HSP70 |
| CLIBASIA_05610 | 44 | | putative phage terminase, large subunit | NA | Terminase_6N/C |
| CLIBASIA_05635 | 40 | | hypothetical protein | NA | NA |
| **CLIBASIA_01650** | 38 | | isoleucine--tRNA synthetase | Aminoacyl-tRNA synthetases | tRNA-synt_1 |
| CLIBASIA_03715 | 32 | | co-chaperone GroES | RNA degradation | Cpn10 |
| CLIBASIA_01685 | 30 | | Bax inhibitor-1/YccA family protein | NA | Bax1-I |
| **CLIBASIA_01705** | 29 | rpsA | 30S ribosomal protein S1 | Ribosome | S1 |
| CLIBASIA_02540 | 28 | | Hsp20 family protein | RNA degradation | HSP20 |
| CLIBASIA_00965 | 28 | | lytic murein transglycosylase | NA | SLT2 |
| CLIBASIA_03035 | 28 | | ferritin | Enzyme | Ferritin |
| **CLIBASIA_00110** | 27 | rpoB | DNA-directed RNA polymerase subunit beta | RNA polymerase | RNA_pol_Rpb2 |
| CLIBASIA_03115 | 27 | | Flp family type IVb pilin | Secretion system | Flp_Fap |
| **CLIBASIA_00325** | 25 | gyrA | DNA gyrase subunit A | DNA repair & recombination | DNA_gyraseA_C |
| CLIBASIA_00995 | 24 | | porin | NA | Porin_2 domain |
| **CLIBASIA_01945** | 24 | | ribonucleoside-diphosphate reductase subunit alpha | purine/pyrimidine metabolism | Ribonuc_red_lgC/N |
| CLIBASIA_01040 | 22 | | ATP/ADP exchange transporter | Transporter | TLC |
| **CLIBASIA_03375** | 20 | pnp | polyribonucleotide nucleotidyltransferase | RNA degredation | Rnase_PH |
| **CLIBASIA_00105** | 20 | rpoC | DNA-directed RNA polymerase subunit beta' | RNA polymerase | RNA_pol_Rpb1 |

| | | | | | |
|---|---|---|---|---|---|
| CLIBASIA_00745 | 19 | fusA | elongation factor G | Translation factor | GTP_EFTU |
| **CLIBASIA_04415** | 19 | | LysE family translocator | Transporter | LysE |
| CLIBASIA_05585 | 18 | | hypothetical protein | NA | Phage_caspid_2 |
| **CLIBASIA_05620** | 17 | | transposase | NA | HTH_tnp_1 |
| **CLIBASIA_01730** | 15 | fabB | beta-ketoacyl-ACP synthase I | Fatty acid metabolism & biosynthesis | ketoacyl-synt |
| CLIBASIA_01515 | 15 | | 50S ribosomal protein L25/general stress proteinCtc | Ribosome | Ribosomal_TL5_C |
| **CLIBASIA_00870** | 15 | rpoD | RNA polymerase sigma factor RpoD | Flagellar assembly | Sigma70 |
| **CLIBASIA_01005** | 15 | | phenylalanine--tRNA ligase subunit beta | Aminoacyl-tRNA synthetases | B3_4 |
| **CLIBASIA_01070** | 14 | rpsF | 30S ribosomal protein S6 | Ribosome | Ribosomal_S6 |
| **CLIBASIA_03870** | 14 | clpB | ATP-dependent chaperone ClpB | Chaperones and folding catalysts | AAA_2 |
| CLIBASIA_03110 | 14 | | Flp family type IVb pilin | Secretion system | Flp_Fap |
| CLIBASIA_03230 | 13 | | hypothetical protein | NA | FAM184 |
| CLIBASIA_00265 | 13 | | amino acid ABC transporter substrate-binding protein | ABC transporter | SBP_bac_3 |
| CLIBASIA_04060 | 13 | | cold-shock protein | NA | CSD |
| CLIBASIA_05325 | 13 | | SurA N-terminal domain-containing protein | Chaperones and folding catalysts | SurA_N_3 |
| CLIBASIA_03930 | 12 | quaB | IMP dehydrogenase | Purine metabolism | IMPDH |
| **CLIBASIA_04160** | 12 | | valine--tRNA ligase | Aminoacyl-tRNA synthetases | tRNA-synt_1 |
| CLIBASIA_00845 | 12 | | hypothetical protein | NA | NA |
| **CLIBASIA_00780** | 12 | clpX | ATP-dependent Clp protease ATP-binding subunit ClpX | Chaperones and folding catalysts | AAA_2 |
| **CLIBASIA_05365** | 12 | | SDR family oxidoreductase | Fatty acid metabolism & biosynthesis | adh_short-C2 |
| **CLIBASIA_03950** | 12 | | response regulator transcription factor | Two-component system | Response_reg |
| CLIBASIA_05150 | 11 | | hypothetical protein | Transporter | OMP_b-brl |
| **CLIBASIA_00960** | 11 | | aspartate-semialdehyde dehydrogenase | Biosynthesis of amino acids | Semialhyde_dhC |

| | | | | | |
|---|---|---|---|---|---|
| CLIBASIA_04765 | 11 | | 2-oxoglutarate dehydrogenase E1 component | TCA cycle | E1_dh |
| **CLIBASIA_02100** | 11 | fabF | beta-ketoacyl-ACP synthase II | Fatty acid metabolism & biosynthesis | ketoacyl-synt |
| CLIBASIA_03390 | 11 | infB | translation initiation factor IF-2 | Translation factor | IF-2 |
| CLIBASIA_03170 | 11 | sppA | signal peptide peptidase SppA | Peptidase and inhibitor | Peptidase_S49 |
| CLIBASIA_04155 | 11 | | DUF2497 domain-containing protein | NA | DUF2497 |
| CLIBASIA_01540 | 10 | metK | methionine adenosyltransferase | Cysteine and methionine metabolism | S-AdoMet |
| **CLIBASIA_00505** | 10 | | methionine--tRNA ligase | Aminoacyl-tRNA synthetases | tRNA-synt_1 |
| **CLIBASIA_01075** | 10 | | 30S ribosomal protein S18 | Ribosome | Ribosomal_S18 |
| CLIBASIA_01645 | 10 | | helix-turn-helix transcriptional regulator | NA | Peptidase_S24 |
| CLIBASIA_03345 | 10 | tsf | translation elongation factor Ts | Translation factor | EF_TS |
| CLIBASIA_00285 | 10 | | ETC complex I subunit | NA | ETC_C1_NDUFA4 |
| CLIBASIA_01790 | 10 | cyoB | cytochrome o ubiquinol oxidase subunit I | Oxidative phosphorylation | COX1 |
| CLIBASIA_02115 | 10 | fabD | ACP S-malonyltransferase | Fatty acid metabolism & biosynthesis | Acyl_transf_1 |
| CLIBASIA_01935 | 10 | bamE | outer membrane protein assembly factor BamE | NA | SmpA_OmlA |
| CLIBASIA_00015 | 10 | | DUF2800 domain-containing protein | NA | DUF2800 |
| CLIBASIA_02310 | 10 | | EamA family transporter | Transporter | EamA |
| **CLIBASIA_02570** | 10 | dnaA | chromosomal replication initiator protein DnaA | Two-component system | Bac_DnaA |
| CLIBASIA_05565 | 10 | | hypothetical protein | NA | NA |
| **CLIBASIA_03395** | 10 | nusA | transcription termination/antitermination protein NusA | Transcription machinery | NusA_N |
| CLIBASIA_05580 | 10 | | hypothetical protein | NA | NA |
| CLIBASIA_04180 | 10 | | DNA-directed RNA polymerase subunit omega | RNA polymerase | RNA_pol_Rpb6 |
| CLIBASIA_00005 | 10 | | hypothetical protein | NA | Protoglobin |
| CLIBASIA_00150 | 10 | tuf | elongation factor Tu | Translation factor | GTP_EFTU |

| | | | | | |
|---|---|---|---|---|---|
| **CLIBASIA_04800** | 9 | | lysozyme | Enzyme | Phage_lysozyme |
| **CLIBASIA_00590** | 9 | typA | translational GTPase TypA | NA | GTP_EFTU |
| **CLIBASIA_02565** | 9 | rpsT | 30S ribosomal protein S20 | Ribosome | Ribosomal_S20p |
| **CLIBASIA_03655** | 9 | | SAM-dependent DNA methyltransferase | Enzyme | N6_Mtase |
| **CLIBASIA_00890** | 9 | rpsI | 30S ribosomal protein S9 | Ribosome | Ribosomal_S9 |
| CLIBASIA_01510 | 9 | | CarD family transcriptional regulator | Transcription factor | CarD_CdnL_TRCF |
| **CLIBASIA_00885** | 9 | rplM | 50S ribosomal protein L13 | Ribosome | Ribosomal_L13 |
| **CLIBASIA_01620** | 9 | carB | carbamoyl-phosphate synthase large subunit | Pyrimidine metabolism | CPSase_L_D2 |
| **CLIBASIA_02945** | 9 | grpE | nucleotide exchange factor GrpE | Chaperones and folding catalysts | GrpE |
| **CLIBASIA_00080** | 9 | | NADP-dependent malic enzyme | Pyruvate metabolism | PTA_PTB |
| **CLIBASIA_03560** | 9 | | bifunctional folylpolyglutamate synthase/dihydrofolate synthase | Folate biosynthesis | Mur_ligase_M |
| CLIBASIA_05630 | 9 | | hypothetical protein | NA | FMP23 |
| CLIBASIA_RS05615 | 9 | ssrA | ssRA-binding protein | NA | SmpB |
| **CLIBASIA_00270** | 8 | | ABC transporter permease subunit | ABC transporter | BPD_transp_1 |
| **CLIBASIA_00775** | 8 | lon | endopeptidase La | Enzyme | Lon_C |
| **CLIBASIA_04135** | 8 | | histidine--tRNA ligase | Aminoacyl-tRNA synthetases | tRNA-synt_His |
| **CLIBASIA_00135** | 8 | nusG | transcription termination/antitermination factorNusG | Transcription machinery | NusG |
| CLIBASIA_00915 | 8 | | EamA family transporter | NA | EamA |
| CLIBASIA_04755 | 8 | sucC | ADP-forming succinate--CoA ligase subunit beta | TCA cycle | ATP-grasp_2 |
| CLIBASIA_05600 | 8 | | head-tail joining protein | NA | Head-tail_con |
| **CLIBASIA_01520** | 8 | aspS | aspartate--tRNA ligase | Aminoacyl-tRNA synthetases | tRNA-synt_2 |
| CLIBASIA_03820 | 8 | | ribonuclease J | RNA degradation | Lactamase_B_2 |
| **CLIBASIA_01175** | 8 | | L,D-transpeptidase family protein | Peptidase and inhibitor | YkuD |

| | | | | | |
|---|---|---|---|---|---|
| **CLIBASIA_01170** | 8 | glyA | serine hydroxymethyltransferase | Glycine, serine, and threonine metabolism | SHMT |

[a] bolded genes are significant genes from the GO Enrichment

[b] NC = normalized read counts as calculated using the counts function of DESeq2 v.1.30.1

[c] domain analysis gene product was based on the information of KEGG database

**Supplemental Table 3.** Comparison of expression profiles determined by RNA-seq and qRT-qPCR for the 27 predicted core Sec-dependent effectors.

| SDE ID (Thapa et al. 2020)[a] | SDE ID (RNA-seq) | Log2 FC (RNA-seq)[b] | Log2 FC (Thapa et al. 2020)[c] | Log2 FC (Yan et al. 2020)[d] | average Las reads in citrus (RNA-seq)[e] | average Las reads in ACP (RNA-seq)[e] | RT-qPCR citrus[f] |
|---|---|---|---|---|---|---|---|
| CLIBASIA_00185 | CLIBASIA_RS00170 | 1.60 | ND | n/a | 0 | 14.75 | no |
| CLIBASIA_00460 | CLIBASIA_RS00440 | -2.29 | -2.01 | n/a | 0.17 | 0 | yes |
| CLIBASIA_00520 | CLIBASIA_RS00480 | -1.80 | -1.15 | -1.31 | 0.50 | 343.25 | yes |
| CLIBASIA_00525 | CLIBASIA_RS00485 | -1.81 | n/a | -2.40 | 0 | 146 | yes |
| CLIBASIA_00530 | CLIBASIA_RS00490 | -1.05 | -1.23 | -0.84 | 0 | 63.75 | yes |
| CLIBASIA_01000 | not annotated | n/a | ND | n/a | 0 | 0 | no |
| CLIBASIA_02075 | CLIBASIA_RS03390 | -1.14 | psyllid only | n/a | 0.17 | 115 | no |
| CLIBASIA_02145 | CLIBASIA_RS03320 | -1.22 | 7.34 | 1.86 | 0.17 | 148.75 | yes |

| | | | | | | | |
|---|---|---|---|---|---|---|---|
| CLIBASIA_03120 | not annotated | n/a | psyllid only | n/a | 0 | 0 | no |
| CLIBASIA_03160 | CLIBASIA_RS02325 | 1.07 | 0.26 | n/a | 2.5 | 292.25 | yes |
| CLIBASIA_03230 | CLIBASIA_RS02250 | 0.38 | 1.20 | 3.52 | 8.17 | 1282 | yes |
| CLIBASIA_03295 | CLIBASIA_RS02190 | -1.72 | -1.15 | n/a | 0 | 133.75 | yes |
| CLIBASIA_04025 | CLIBASIA_RS03990 | 1.95 | 2.55 | 3.40 | 0.17 | 15.25 | yes |
| CLIBASIA_04030 | CLIBASIA_RS03995 | 1.10 | 0.88 | 0.83 | 0 | 29 | yes |
| CLIBASIA_04040 | CLIBASIA_RS04000 | -0.76 | 0.36 | 2.43 | 0 | 87.5 | yes |
| CLIBASIA_04055 | CLIBASIA_RS04015 | 0.45 | ND | n/a | 0.17 | 40.5 | no |
| CLIBASIA_04320 | CLIBASIA_RS04265 | -0.21 | -1.89 | 4.5 | 3.17 | 728.75 | yes |
| CLIBASIA_04405 | CLIBASIA_RS04345 | 3.17 | ND | 2.86 | 1.33 | 32.25 | no |
| CLIBASIA_04410 | CLIBASIA_RS04350 | -1.95* | ND | n/a | 1.50 | 924.25 | yes |
| CLIBASIA_04530 | CLIBASIA_RS04460 | 0.69 | n/a | 2.67 | 0 | 21.5 | no |
| CLIBASIA_04560 | CLIBASIA_RS04490 | 1.46 | n/a | 2.04 | 1.00 | 63.25 | yes |
| CLIBASIA_04580 | CLIBASIA_RS04510 | 0.51 | n/a | 2.04 | 1.50 | 168 | yes |
| CLIBASIA_04690 | CLIBASIA_RS04610 | -0.16 | ND | n/a | 0 | 30.5 | no |
| CLIBASIA_04970 | not annotated | n/a | ND | n/a | 0 | 0 | no |
| CLIBASIA_05315 | CLIBASIA_RS05165 | 4.91* | 3.16 | n/a | 3.33 | 24.5 | yes |
| CLIBASIA_05320 | CLIBASIA_RS05170 | 2.41 | 4.62 | 4.39 | 3.17 | 100.25 | yes |

| | | | | | | | |
|---|---|---|---|---|---|---|---|
| CLIBASIA_05330 | CLIBASIA_RS05180 | -0.91 | ND | 1.51 | 0.17 | 111.5 | yes |

[a] bolded genes belong to the updated list of SDE candidates

[b] $\log_2$ fold change ($\log_2$FC) values in Las gene expression in grapefruit samples compared with insect samples for putative Sec-dependent effectors (SDEs) from our RNA-seq study where positive values indicate higher expression in citrus host, negative values indicate higher expression in insect host, and 'n/a' indicates not available. * indicates significant DEG.

[c] $\log_2$ fold change ($\log_2$FC) values in Las gene expression in citrus samples compared with insect samples by qRT-PCR for predicted SDEs from Thapa et al. (2020). Positive values indicate higher expression in citrus host, negative values indicate higher expression in insect host, 'psyllid only' indicates expression in only insect host, 'ND' indicates not detected, and n/a not available.

[d] $\log_2$ fold change ($\log_2$FC) values in Las gene expression in citrus samples compared with insect samples by qRT-PCR for predicted SDEs from Yan et al. (2013). Positive values indicate higher expression in the citrus host, negative values indicate higher expression in the insect host, and n/a indicates not available.

[e] values indicate the average number of reads that mapped to each putative Las SDE from all bacterial samples isolated from grapefruit and non-enriched ACP respectively. n/a indicates not available.

[f] expression of putative SDEs in citrus hosts detected by qRT-PCR. yes: significant ΔCt citrus=(Ct av Las gene in healthy citrus- Ct av Las gene in Las infected citrus); no: non-significant ΔCt citrus.
